# LoCoFlow: Local-Context Flow Matching for Single-Cell Dynamics and Fate Inference

**DOI:** 10.64898/2026.10.15.758594

**Authors:** Chang Liu, Zibo Guan, Xun Lan

## Abstract

Single-cell sequencing profiles cellular states at discrete time points, providing population-level snapshots from which developmental and disease-associated trajectories can be reconstructed. Existing optimal-transport and flow-matching approaches typically infer temporal couplings at the level of individual cells, with limited representation of the local manifold geometry of cell-state space and the uncertainty inherent in lineage diversification. We introduce LOCOFLOW, a flow-matching framework that combines local point-cloud geometry with conditional fate uncertainty. Overlapping patches define hierarchical transport between populations, and an endpoint-preserving neural bridge regularizes motion and deformation within these patches. Bridge velocities supervise a conditional field with patch-shared and cell-specific latent variables, from which alternative trajectories can be sampled without future observations. Temporal and spatial benchmarks show improvements in transport distance. On lineage-traced LARRY data, LOCOFLOW reduces mean raw Brier error from 0.7990 to 0.6620 relative to scDiffEq at the same number of endpoint draws. Expression perturbations further reveal distinct roles for focal-cell and neighborhood information: geometric-patch models show aligned background and whole-patch responses, whereas random-minibatch models show opposing focal and background responses. These findings support local context as a useful representation for probabilistic fate inference and provide testable hypotheses about how it shapes predicted responses.

## 1 Introduction

Differentiation during embryonic development, tissue regeneration, and the adaptation of tumor cells to therapy all involve changes in cell states over time. While single-cell sequencing and spatial transcriptomics can obtain the molecular states and tissue locations associated with these processes, the measurements are typically destructive so that data obtained at different time points represent snapshots of distinct cell populations, leaving the true longitudinal correspondences and intermediate states unobserved. Additionally, reconstructing continuous trajectories from snapshots alone yields multiple possible solutions, necessitating the use of temporal, geometric, or biological constraints to guide inference (Zhang et al., 2026; Weinreb et al., 2018). Furthermore, lineage tracing studies have demonstrated that cells with similar transcriptional states can exhibit divergent fate tendencies, suggesting that a single cell’s instantaneous expression vector is insufficient to fully characterize its subsequent changes (Weinreb et al., 2020).

Existing methods have progressively advanced trajectory inference from cell ordering to continuous dynamic modeling. Pseudotime reconstructs the temporal sequence of states, while RNA velocity indicates local directions of change; optimal transport connects cell populations across different time points, and continuous flow models enable the inference of states at unsampled time points (Trapnell et al., 2014; La Manno et al., 2018; Schiebinger et al., 2019; Tong et al., 2020). In recent years, single-cell trajectory modeling has evolved from point-level matching to continuous-time frameworks that simultaneously account for data manifold geometry, stochastic dynamics, and shifts in population distributions. Geometric constraints align inferred paths more closely with observable cell states; probabilistic transport preserves multiple potential temporal correspondences; and distribution-level flow matching enables the treatment of point clouds themselves as evolving entities. While these advances lay the foundation for reconstructing complex cell trajectories, a key challenge remains: how to represent the local neighborhood of an anchored cell as a whole that evolves over time, while comprehensively characterizing its global motion, internal structural changes, and matching uncertainties. (Huguet et al., 2022; Kapuśniak et al., 2024; Yu et al., 2026; Klein et al., 2024; Haviv et al., 2025).

To address this, we introduce LOCOFLOW. For each anchor cell, LOCOFLOW constructs a point cloud from its local neighborhood, thereby extending the fundamental unit of dynamic modeling from an isolated expression vector to a local population state. First, matching point cloud distributions across time points integrates cell states with neighborhood geometry and employs probabilistic pairing to preserve potential multiple successor correspondences. Second, LOCOFLOW decomposes point cloud evolution into the movement of items and changes in their internal discrete structure: the former characterizes local motion as conforming to the manifold, while the latter describes neighborhood variations—such as expansion, contraction, and heterogeneity. Finally, point cloud itself provides a distributional representation for studying fate bifurcation. Its biological significance requires validation through lineage tracing, held-out time-point prediction, or perturbation data.

## 2 Related Work

### From cell ordering to continuous-time dynamics

Pseudotime methods like Monocle arrange cell states based on expression differences; RNA velocity estimates the direction of local change using unspliced and spliced RNA; and CellRank integrates directional information with cell-to-cell similarity to calculate the probabilities of cells reaching various terminal states. While these methods provide information on sequence, direction, or fate propensity, they do not directly observe cell correspondences across distinct experimental time points (Trapnell et al., 2014; La Manno et al., 2018; Lange et al., 2022). For time-series snapshots, PRESCIENT learns a stochastic differentiation landscape to predict the outcomes of interventions; scNODE employs neural ordinary differential equations in latent space to generate expression states at unobserved time points; and scDiffEq uses neural stochastic differential equations to simultaneously model directed changes and stochastic diffusion (Yeo et al., 2021; Zhang et al., 2024; Vinyard et al., 2025).

### Transport-based temporal matching

Waddington-OT employs optimal transport to infer ancestor-descendant relationships between cell populations at different time points during reprogramming, while TrajectoryNet utilizes dynamic optimal transport for continuous trajectory generation. To account for changes in cell numbers over time, TIGON jointly reconstructs state movement and population growth, and DeepRUOT further incorporates stochastic dynamics as well as mass changes resulting from proliferation and death; meanwhile, moscot applies transport methods to large-scale spatiotemporal and multimodal single-cell data (Schiebinger et al., 2019; Tong et al., 2020; Sha et al., 2024; Zhang et al., 2025; Klein et al., 2025). These methods demonstrate that temporal matching requires the simultaneous consideration of state distances, population distributions, and non-conservative changes inherent in biological processes.

### Geometric constraints and distribution-level flow matching

Cellular expression states often concentrate on low-dimensional spaces; consequently, direct interpolation in Euclidean space may not align with observed state transitions. MIOFlow learns population trajectories in a latent space that preserves manifold distances; Metric Flow Matching (MFM) constructs approximate geodesic interpolations using a data-induced Riemannian metric; PACE employs a state- and time-dependent anisotropic metric to reduce the cost of movement along local tangents while penalizing movement in the normal direction (Huguet et al., 2022; Kapuśniak et al., 2024; Yu et al., 2026). Flow matching trains continuous models by learning time-varying velocity fields, typically avoiding the need to simulate full trajectories during each parameter update. SF^2^M integrates flow matching and score matching within the Schrödinger bridge framework to characterize stochastic transitions; GENOT combines entropic regularized transport with flow matching to learn probabilistic mappings; VGFM jointly learns cellular state dynamics and population growth (Tong et al., 2024b; Klein et al., 2024; Wang et al., 2025). Wasserstein Flow Matching further demonstrates that empirical point clouds can themselves serve as distributions, allowing the formulation of flow-matching objectives within the space of distributions (Haviv et al., 2025). LOCOFLOW relates to these works by simultaneously leveraging transport, local geometry, and point-cloud representations; its specific focus lies in how local populations—defined by anchored cell neighborhoods—match across time, and how the movement of their centers combined with changes in their internal structures characterizes cellular trajectories.

## 3 Method

### Problem and design

We observe cell-state clouds 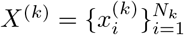 at times 0 = *t*_0_ *<* · · · *< t*_K_, with 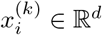 and empirical distributions 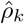. The task is to infer dynamics between snapshots and predict a distribution of future states for each starting cell. Matching the marginals 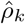 alone does not determine these conditional outcomes. LOCOFLOW uses local point-cloud patches to constrain correspondence and path geometry, while retaining stochastic variation in the inferred dynamics.

Training has two stages (Figure 1). Stage 1 constructs hierarchical transport couplings and learns endpoint-preserving, geometry-regularized bridges. Stage 2 distills their velocities into a conditional field with patch-shared and cell-specific latent variables. At inference, latent codes are drawn without future observations and held fixed during each interval. This separates uncertainty about outcomes from the geometric constraints on the paths leading to them.

**Figure 1.**
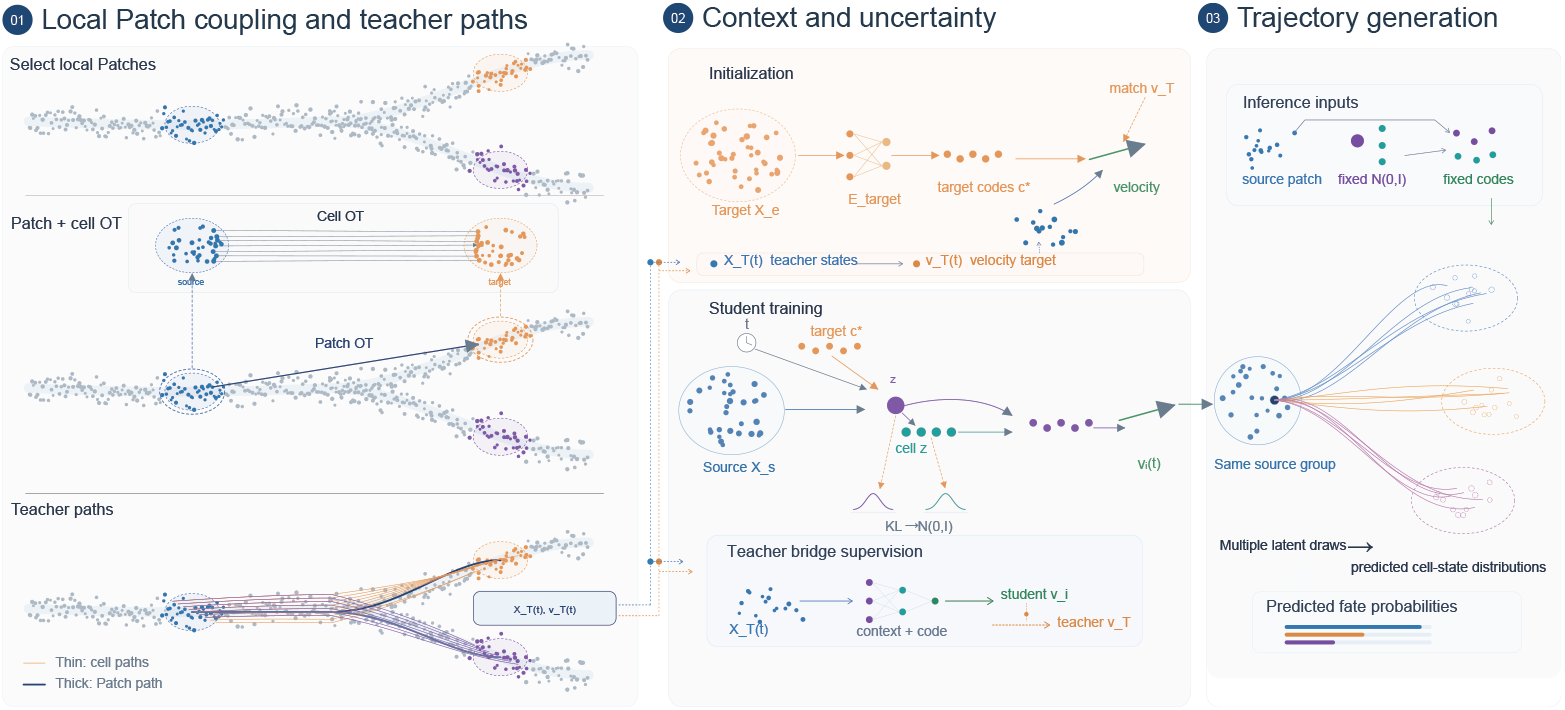
Local geometry and conditional dynamics. Hierarchical transport couples overlapping patches; Stage 1 learns geometry-regularized bridges; Stage 2 distills their velocities into a hierarchical latent-conditioned field. Targets supervise training only. At inference, prior latent draws generate codes and predicted states propagate between intervals.

### 3.1 Local PATCHES AND HIERARCHICAL CORRESPONDENCE

For expression data, patches are geodesic neighborhoods on a nearest-neighbor graph, with support size selected by a local low-rank residual. For spatial data, they are fixed-size neighborhoods in measured coordinates. Patches overlap, but membership weights partition each cell’s mass:

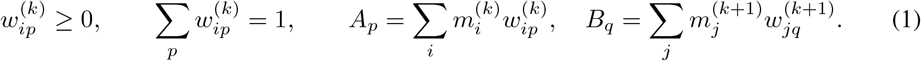

Here 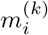 is the cell mass used for transport, and *A*_*p*_, *B*_*q*_ are the resulting source and target patch masses. The balanced problem below requires equal total source and target mass. With patch centers *µ*_p_, *µ*_*q*_ and low-rank covariances Σ_*p*_, Σ_*q*_, the patch-level plan solves

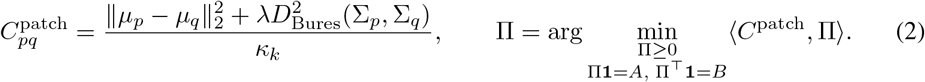

The Bures term compares patch dispersion as well as location; *κ*_*k*_ sets the interval’s cost scale. Computation uses a restricted candidate support and averages feasible plans under multiple cost perturbations.

Within each retained patch pair, an entropy-regularized cell-level plan *T* ^pq^ uses squared-distance cost divided by *κ*_*k*_ and marginals 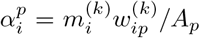 and 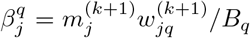. The full local objective is given in Appendix A. Aggregating these plans gives

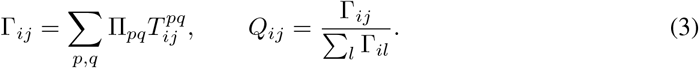

Membership normalization prevents overlap from duplicating transport mass (Appendix A). *Q* defines a conditional distribution of target cells for each source, rather than a single hard assignment. It supplies paired endpoint sets with patch context, 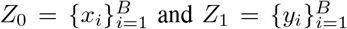, for bridge training.

### 3.2 Stage 1: GEOMETRY-REGULARIZED BRIDGE TRANSPORT

Endpoint correspondence does not specify the intermediate path. For Δ*t*_*k*_ = *t*_*k*+1_ − *t*_*k*_ and local phase *τ* = (*t* − *t*_*k*_)*/*Δ*t*_*k*_, we learn

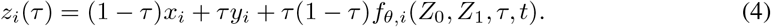

The bridge combines cell features with patch-mean features in a permutation-equivariant network. The residual prefactor enforces both endpoints exactly while allowing curved paths. Differentiation gives

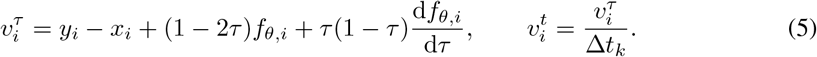

Automatic differentiation includes the dependence of the time input *t* on *τ*. The bridge objective is

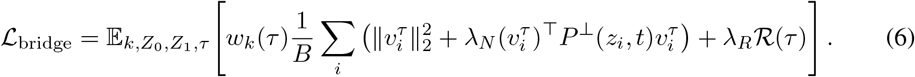

Kinetic energy discourages unnecessary motion. The precomputed local normal field *P*^⊥^ penalizes velocity orthogonal to the estimated local geometry. The radial term *R* compares the instantaneous rate of change of source-neighbor distances with their endpoint distance difference (Appendix B). It constrains relative deformation, not common translation. The normalized phase weight *w*_k_ depends on the change in population radius between endpoints. These terms are geometric priors on unobserved paths, not observations of an intermediate manifold.

### 3.3 Stage 2: CONDITIONAL VELOCITY DISTILLATION

The bridge uses both endpoints and therefore cannot be used directly when the future is unknown. We distill it into a conditional field *V*_*ω*_(*Z, u* | *Z*_0_, *C, k*), where *Z* is the evolving patch, *Z*_0_ its source context, *C* a set of conditioning codes, and *k* the interval. Global time is normalized as *u* = (*t* − *t*_0_)*/*(*t*_*K*_ − *t*_0_); the velocity target is consequently 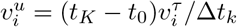.

#### Learning a target representation

With the bridge frozen, a target encoder produces 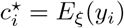. The encoder and velocity network are first trained jointly to match bridge velocities conditional on these codes. This initialization makes the codes relevant to the dynamics, rather than assigning them a prescribed fate label. We then freeze the encoder and initialize the next phase from the learned velocity network (Appendix C).

#### Generating codes without targets

At inference, the target codes are unavailable. A generator *c*_*i*_ = *D*_*ψ*_(*Z*_0_, *η*_*P*_, *η*_*i*_, *k*) instead combines source context with a patch-shared latent *η*_P_ and a cell-specific latent *η*_*i*_. Their joint prior is *p*(*η*) = *N*(*η*_*P*_ ; 0, *I*) Π*i* (*η*_*i*_; 0, *I*). Training uses a hierarchical Gaussian posterior

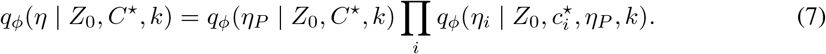

The patch posterior receives the mean and standard deviation of standardized target codes; each cell posterior receives its own code. Reparameterized latent samples generate 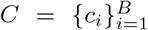. Writing 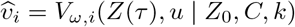, training minimizes

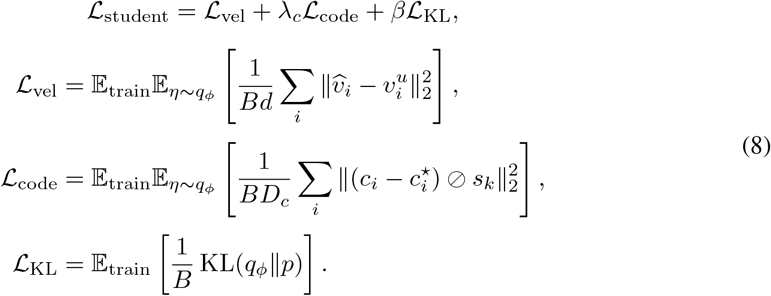

Here *D*_*c*_ is the code dimension and *s*_*k*_ the interval-specific training-code standard deviation, floored componentwise at 0.05; ⊘ denotes elementwise division. The terms respectively fit velocities, reconstruct target codes, and align the training posterior with the inference prior. Dividing the joint KL by *B* allocates the shared-latent penalty across the patch. The shared latent can correlate outcomes within a population, while cell latents retain individual variation.

#### Inference

At each interval’s start, we form source patches, draw both latent levels from the prior, and generate codes. Holding source context and codes fixed, we integrate

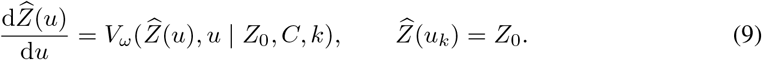

Predicted states become the source context for the next interval, where new latents are drawn. No intermediate observation resets the trajectory. Repeated latent draws yield a conditional distribution of outcomes; each draw defines an ODE trajectory rather than continuous-time diffusion.

## 4 Experiments

The experiment evaluates Locoflow across three aspects. First, we assess whether Locoflow improves the performance of held-out reconstruction for single-cell differentiation and reprogramming processes based on destructive snapshots, and whether it remains effective as the dimensionality of the representation increases. Second, we examine whether Locoflow can predict fate uncertainty during cell differentiation and identify key factors governing fate decisions through perturbation experiments. Third, we evaluate whether Locoflow improves held-out reconstruction performance and distinguishes between environmental and intrinsic influences based on using spatiotemporal transcriptomic data.

### 4.1 Hold OUT TIME POINT RECONSTRUCTION

We evaluate held-out population reconstruction on embryoid-body (EB) differentiation, iPSC reprogramming (Liu et al., 2020), and *C. elegans* development. EB is tested in 2D and PCA50; iPSC in PCA10 and PCA50, holding out days 4, 8, 12, and 16 separately; and *C. elegans* under five-fold temporal holdout. Test time points are excluded from fitted preprocessing and neural training. Dataset-specific variants appear in Table 1. We compare methods in the same representation using empirical W1 and W2 distances and MMD (Appendix D); reported uncertainty is the sample standard deviation across training seeds.

**Table 1:** Time-held-out population prediction. Values are means ± training-seed sample SD; lower is better. Bold and underlining mark the best and second-best means in each setting, determined before rounding.

| Dataset | Method | W1 ↓ | W2 ↓ | MMD ↓ |
| --- | --- | --- | --- | --- |
| EB-2D | OT-CFM | 0.2831 ± 0.0118 | 0.3670 ± 0.0109 | 0.0977 ± 0.0066 |
| | MFM | 0.3540 ± 0.0503 | 0.4459 ± 0.0562 | 0.1304 ± 0.0372 |
|  | PACE | **0.2443 ± 0.0109** | **0.3196 ± 0.0100** | **0.0957 ± 0.0075** |
|  | LoCoFlow | <u>0.2645 ± 0.0177</u> | <u>0.3468 ± 0.0268</u> | <u>0.0957 ± 0.0111</u> |
| EB-PCA50 | OT-CFM | <u>7.3485 ± 0.0205</u> | <u>7.4855 ± 0.0212</u> | **0.1187 ± 0.0013** |
| | MFM | 7.4055 ± 0.0214 | 7.5440 ± 0.0222 | 0.1197 ± 0.0010 |
| | PACE | 7.3509 ± 0.0218 | 7.4884 ± 0.0218 | 0.1187 ± 0.0022 |
| | LoCoFlow | **6.4635 ± 0.0064** | **6.6036 ± 0.0059** | 0.2269 ± 0.0018 |
| iPSC-2D | OT-CFM | 0.5132 ± 0.0135 | 0.5899 ± 0.0130 | **0.4309 ± 0.0089** |
| | MFM | 0.9288 ± 0.0336 | 1.0096 ± 0.0498 | 0.6514 ± 0.0211 |
| | PACE | 0.5003 ± 0.0060 | 0.5702 ± 0.0055 | 0.4321 ± 0.0130 |
|  | LoCoFlow | **0.4455 ± 0.0053** | **0.5314 ± 0.0074** | <u>0.4349 ± 0.0107</u> |
| iPSC-PCA10 | OT-CFM | 2.8317 ± 0.0244 | 3.1407 ± 0.0240 | <u>0.4348 ± 0.0022</u> |
| | MFM | 2.8568 ± 0.0247 | 3.1622 ± 0.0244 | 0.4391 ± 0.0040 |
| | PACE | **2.7674 ± 0.0252** | 3.1002 ± 0.0260 | **0.4286 ± 0.0045** |
| | LoCoFlow | <u>2.7698 ± 0.0226</u> | **3.0832 ± 0.0257** | 0.4767 ± 0.0052 |
| iPSC-PCA50 | OT-CFM | 7.2240 ± 0.0349 | 7.7028 ± 0.0462 | **0.2662 ± 0.0010** |
| | MFM | 7.2286 ± 0.0334 | 7.7151 ± 0.0481 | 0.2676 ± 0.0034 |
| | PACE | 7.2809 ± 0.0224 | 7.8494 ± 0.0454 | 0.2669 ± 0.0011 |
|  | LoCoFlow | **6.5523 ± 0.0128** | **6.9946 ± 0.0126** | <u>0.3442 ± 0.0016</u> |
| *C. elegans* | OT-CFM | 6.9159 ± 0.1207 | 7.0882 ± 0.1633 | 0.2260 ± 0.0095 |
| | MFM | 7.1818 ± 0.5929 | 7.4880 ± 0.7875 | 0.2290 ± 0.0320 |
| | PACE | 6.9182 ± 0.4450 | 7.1621 ± 0.5494 | **0.2183 ± 0.0178** |
| | LoCoFlow | **6.5573 ± 0.0248** | **6.7337 ± 0.0155** | 0.3157 ± 0.0028 |

Locoflow has the lowest mean W2 in five of six settings. Relative to OT-CFM (Tong et al., 2024a), W2 falls by 11.8% on EB-PCA50 (7.4855 to 6.6036) and 9.2% on iPSC-PCA50 (7.7028 to 6.9946); the iPSC-PCA10 gain is smaller, and PACE performs better on EB-2D. However, Locoflow has higher MMD than OT-CFM on EB-PCA50 (0.2269 versus 0.1187) and iPSC-PCA50 (0.3442 versus 0.2662). In the Figures S01–S04, Locoflow’s predictions appear more concentrated in some regions of the cell-state space; such local density discrepancies may partly explain the higher MMD despite lower W2.

### 4.2 Fate Prediction

LARRY links early states to measured clonal descendants (Weinreb et al., 2020), testing conditional fate probabilities rather than a held-out population marginal. Lineage labels are reserved for evaluation. Figure 2 shows the observed landscape; Figure S06 illustrates progenitors with one to three observed differentiated fates.

**Figure 2.**
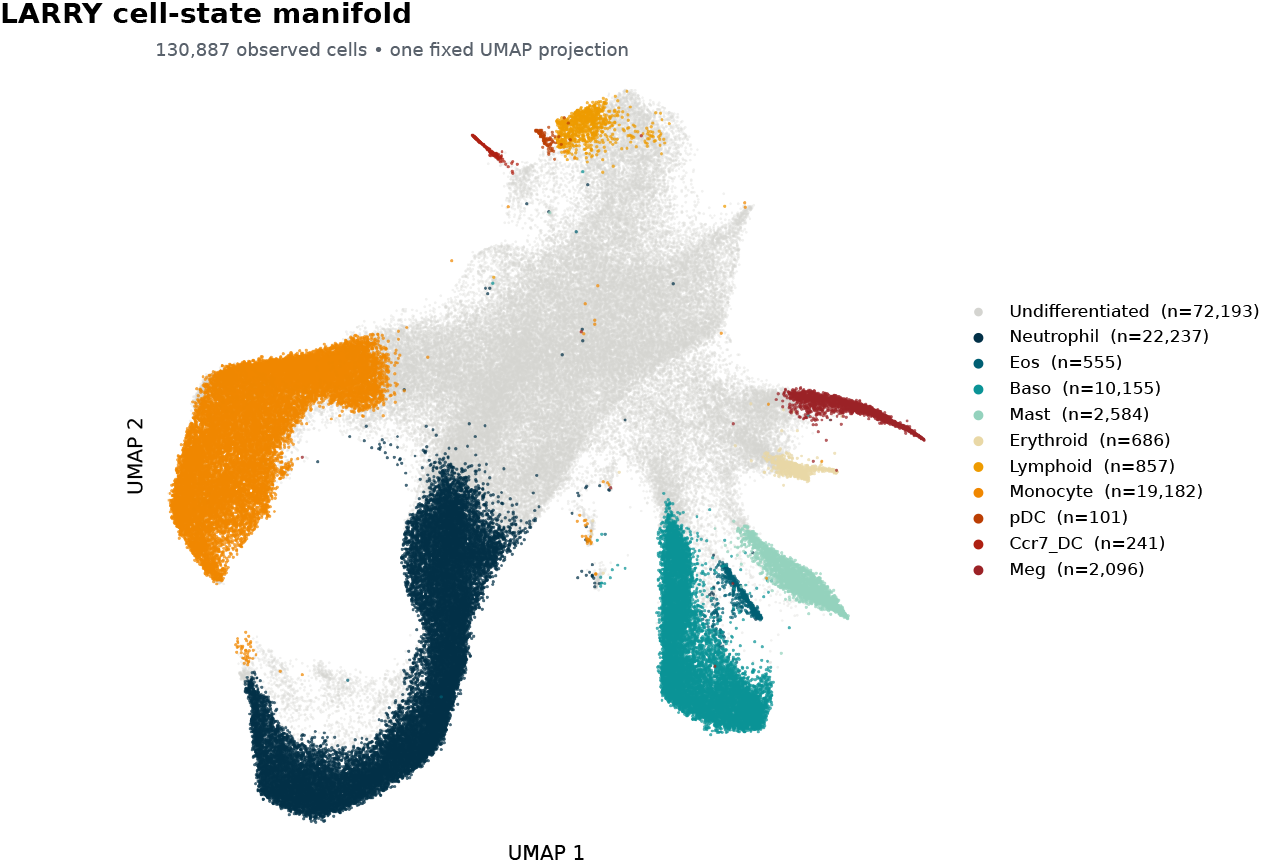
Observed LARRY cell-state landscape. Fixed UMAP of 130,887 observed cells, including undifferentiated cells, colored by annotation. This is neither a predicted trajectory nor a lineage tree.

We distinguish raw from differentiated-conditional fate probabilities, as in the scDiffEq evaluation workflow (Vinyard et al., 2025); the relevant evaluation details are given in Appendix D. Here we use the Brier score, rather than the negative cross entropy in its Figure S7 notebook, to evaluate the raw endpoint vector, including the undifferentiated class U:

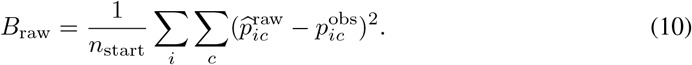

The observed descendant proportion 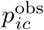 has U set to zero for this task. Dominant differentiated-fate accuracy and balanced accuracy are also reported; all-U predictions remain in evaluation.

At 2,000 endpoint draws per starting cell, Locoflow achieves a raw Brier score of 0.6620 ± 0.0079, lower than that of VRFM, another variational flow-matching method (Table 2). It also improves on scDiffEq. These results favor Locoflow for predicting the full fate-probability vector; they do not imply uniformly better dominant-fate classification.

**Table 2:**
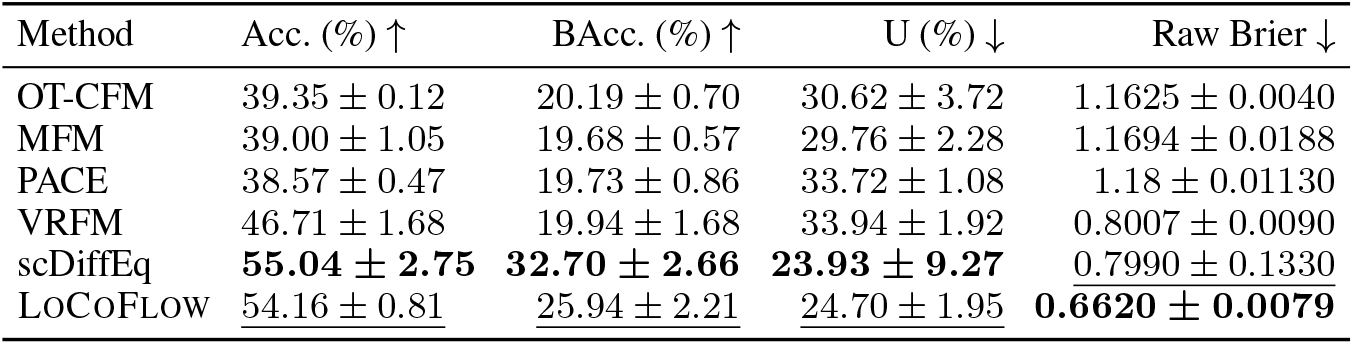
LARRY test-set fate prediction. Values are means ± training-seed sample SD. Draw counts are per starting cell. U is the unfiltered undifferentiated endpoint mass; lower is better in this evaluation. Bold and underlining mark the best and second-best means, respectively.

| Method | Acc. (%) ↑ | BAcc. (%) ↑ | U (%) ↓ | Raw Brier ↓ |
| --- | --- | --- | --- | --- |
| OT-CFM | 39.35 ± 0.12 | 20.19 ± 0.70 | 30.62 ± 3.72 | 1.1625 ± 0.0040 |
| MFM | 39.00 ± 1.05 | 19.68 ± 0.57 | 29.76 ± 2.28 | 1.1694 ± 0.0188 |
| PACE | 38.57 ± 0.47 | 19.73 ± 0.86 | 33.72 ± 1.08 | 1.18 ± 0.01130 |
| VRFM | 46.71 ± 1.68 | 19.94 ± 1.68 | 33.94 ± 1.92 | 0.8007 ± 0.0090 |
| scDiffEq | **55.04 ± 2.75** | **32.70 ± 2.66** | **23.93 ± 9.27** | 0.7990 ± 0.1330 |
| LoCoFlow | <u>54.16 ± 0.81</u> | <u>25.94 ± 2.21</u> | <u>24.70 ± 1.95</u> | **0.6620 ± 0.0079** |

To test whether the learned dynamics predict interpretable responses to changes in gene expression, we performed in silico perturbations of *Cebpe, Dach1, Lmo4*, and *Mxd1*. We jointly clamped their expression over *z* ∈ {−10, *−*7.5, …, 10} in the focal progenitor (target), its surrounding patch cells (background), or both. We then simulated descendant fates and quantified the monocyte and neutrophil fractions and their ratio (Table 3).

**Table 3:** Joint four-gene perturbation in LARRY: *Cebpe, Dach1, Lmo4*, and *Mxd1*. Entries give neutrophil% / monocyte%, with their ratio below.

| $z$ | scDiffEq | Target only | Background only | All members |
| --- | --- | --- | --- | --- |
|  | 11.6/32.4 | 23.9/38.9 | 12.8/51.3 | 12.2/52.0 |
| −10 | (0.3580) | (0.6144) | (0.2495) | (0.2346) |
|  | 13.2/32.7 | 24.2/38.7 | 14.8/48.1 | 13.9/48.9 |
| −7.5 | (0.4037) | (0.6253) | (0.3077) | (0.2843) |
|  | 15.8/31.5 | 24.3/38.4 | 18.3/44.9 | 17.8/45.3 |
| −5 | (0.5016) | (0.6328) | (0.4076) | (0.3929) |
|  | 18.2/30.6 | 24.6/38.2 | 21.4/41.6 | 21.3/41.9 |
| −2.5 | (0.5948) | (0.6440) | (0.5144) | (0.5084) |
|  | 20.9/27.5 | 24.8/38.0 | 24.9/38.0 | 24.9/38.0 |
| 0 | (0.7600) | (0.6526) | (0.6553) | (0.6553) |
|  | 23.2/27.2 | 24.9/38.0 | 28.6/33.6 | 29.0/33.4 |
| 2.5 | (0.8529) | (0.6553) | (0.8512) | (0.8683) |
|  | 27.0/24.2 | 25.0/38.0 | 32.0/30.1 | 32.7/29.5 |
| 5 | (1.1157) | (0.6579) | (1.0631) | (1.1085) |
|  | 35.0/19.0 | 25.8/37.9 | 36.5/26.1 | 38.0/25.5 |
| 7.5 | (1.8421) | (0.6807) | (1.3985) | (1.4902) |
|  | 42.5/14.4 | 26.1/37.4 | 40.7/21.8 | 41.8/21.2 |
| 10 | (2.9514) | (0.6979) | (1.8670) | (1.9717) |

Increasing the clamp from *z* = −10 to 10 in geometrically defined patches shifted predicted fates from monocytes toward neutrophils when applied to the background cells: the respective fractions changed by −29.5 and +27.9 percentage points, and the neutrophil-to-monocyte ratio rose from 0.2495 to 1.8670. Perturbing all patch members produced a similar shift (−30.8 and +29.6 points), whereas perturbing only the focal cell had smaller effects (−1.5 and +2.2 points; Figure S10). Figure 3 compares three progenitors under the *z* = 0 clamp, and Figure S07a shows their responses across perturbation levels. A separate single-gene screen nominates lineage-associated candidate regulators (Figure S08).

**Figure 3.**
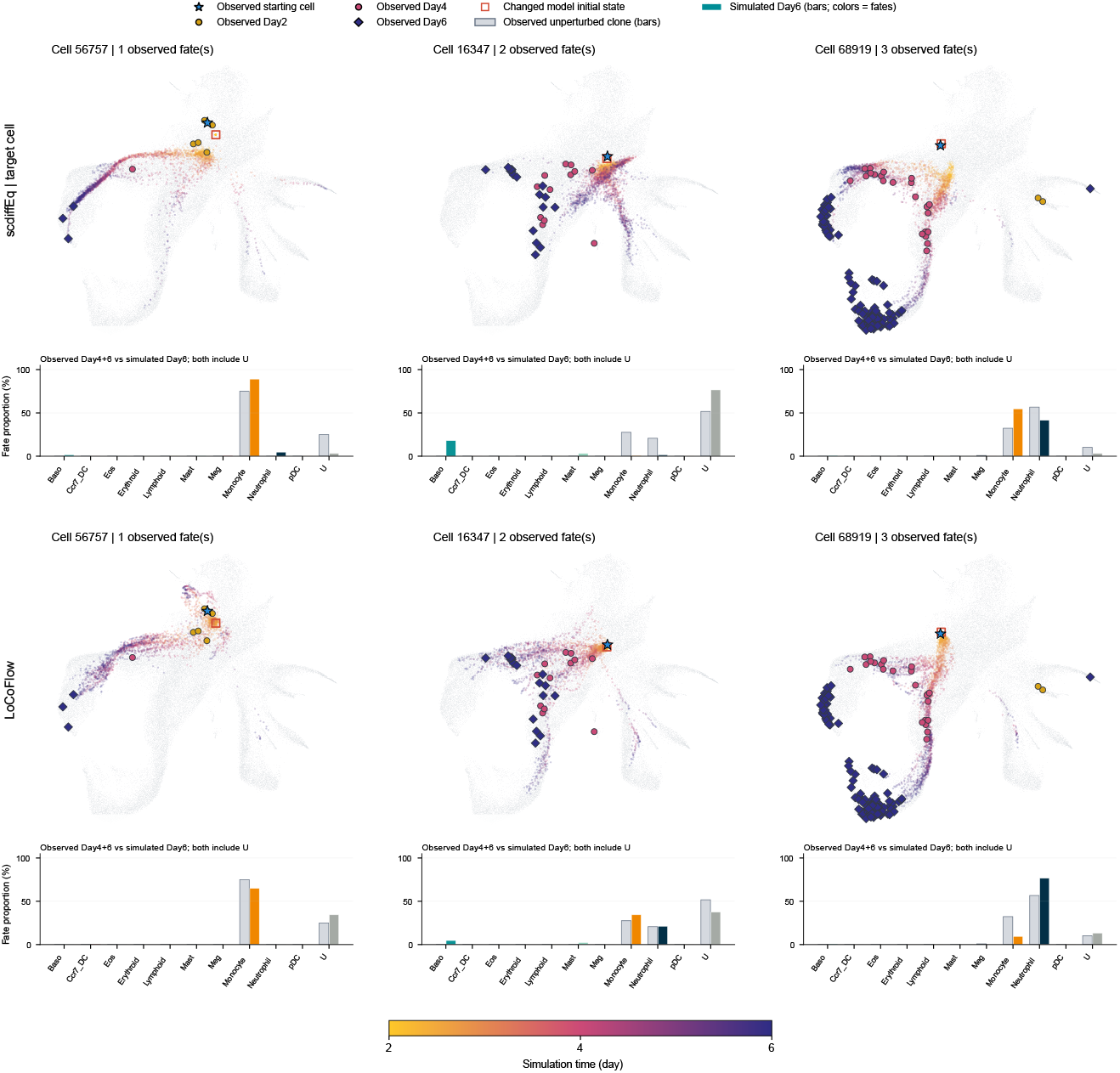
Focal-cell and contextual perturbations at *z* = 0. Columns show progenitors 56757, 16347, and 68919 with one, two, and three observed differentiated fates. Top: scDiffEq target-cell perturbation; bottom: LOCOFLOW target-only perturbation. Background and all-member arms continue below. Points show simulated states; bars compare simulated Day 6 fates with untreated Day 4+6 descendants, retaining U. The joint clamp affects *Cebpe, Dach1, Lmo4*, and *Mxd1*.

Together, these predictions indicate that Locoflow uses local context when assigning progenitor fates. By contrast, randomly assembled patches produced opposing focal-cell and background responses, with greater sensitivity to focal-cell perturbation in that configuration (Figure S09).

Ablations support the role of context and latent variation (Tables 5 and 6): removing neighborhood information yields Brier 0.8464 and 10.00% balanced accuracy; removing hierarchical latents yields Brier 1.0039. Random-minibatch OT gives a lower Brier than the full model (0.6212 versus 0.6833) but lower balanced accuracy (18.24% versus 26.19%).

### 4.3 Axolotl Brain Regeneration Interpolation

For Axolotl Brain Regeneration, we combine measured spatial coordinates with ten principal components computed from 2,000 highly variable genes (HVGs) to represent cell states. Locoflow constructs each patch using only the measured spatial coordinates. We first predict the 5-DPI cell population from 2 DPI; Table 4 aggregates five evaluation subsamples within each of five training seeds.

**Table 4:** Spatial prediction, 2→5 DPI (Axolotl Brain Regeneration). Bold and underlining mark the best and second-best means, respectively.

| Method | W2 ↓ | Unbiased MMD<sup>2</sup> ↓ | Energy ↓ |
| --- | --- | --- | --- |
| OT-CFM | <u>0.6503 ± 0.0042</u> | **0.0127 ± 0.0007** | **0.0535 ± 0.0033** |
| MFM | 0.6608 ± 0.0069 | 0.0334 ± 0.0035 | 0.0837 ± 0.0086 |
| PACE | 0.6562 ± 0.0044 | <u>0.0146 ± 0.0007</u> | <u>0.0615 ± 0.0031</u> |
| ContextFlow | 0.6710 ± 0.0108 | 0.0179 ± 0.0023 | 0.0774 ± 0.0097 |
| LoCoFlow | **0.6128 ± 0.0040** | 0.0182 ± 0.0009 | 0.0629 ± 0.0038 |

Locoflow achieves W2 0.6128 ± 0.0040, 5.8% below OT-CFM’s 0.6503. OT-CFM performs better on unbiased MMD^2^ (0.0127 versus 0.0182) and energy distance (0.0535 versus 0.0629). In the measured-coordinate views (Figure S11), the predicted spatial distribution remains visually similar to that at 2 DPI, suggesting that additional spatial registration or alignment may be needed to capture the observed 5-DPI organization.

We further perturb each HVG individually in either a 10-DPI reaEGC (target) or the non-reaEGC cells in its spatial patch (background). For each perturbed starting cell, we simulate 100 trajectories to 15 DPI and summarize changes in predicted neuronal fate probability across five training seeds. Target and background interventions generally produce similar directional responses, although target interventions tend to have larger effects. Decreasing *SLC1A3* or *GLUL*, and increasing *FKBPB* or *CHGA*, yield particularly pronounced increases in predicted neuronal fate probability (Figure S12).

## 5 Discussion AND Conclusion

LOCOFLOW combines geometry-aware correspondence, regularized bridge paths, and latent-conditioned dynamics to model cell-state transitions. Across differentiation and reprogramming datasets, it improves held-out reconstruction on W1/W2 in most evaluated settings, including higher-dimensional representations. On LARRY, its lower raw Brier score than the variational flow-matching baseline VRFM and scdiffeq indicates more accurate prediction of full fate-probability vectors. However, these gains are not consistent: higher MMD on some datasets and lower balanced accuracy than scDiffEq show that local population structure and less frequent fates remain challenging. In Axolotl Brain Regeneration, spatially defined patches improve W2, although the predicted spatial coordinates remain visually similar to those at 2 DPI, suggesting that additional spatial alignment may be necessary.

The perturbation analyses offer a complementary view of the learned dynamics. With geometric patches, background-only and all-member interventions produce similar fate shifts. With randomly assembled patches, focal-cell and background interventions produce opposing responses, and the model is more sensitive to the focal cell. Thus the role of local context is reflected in the structure of predicted responses as well as in aggregate scores. In the Axolotl experiment, target and background gene perturbations generally have effects in similar directions, with larger responses often observed for target perturbations.

These findings require cautious interpretation. Snapshot data do not uniquely identify cellular histories, and the geometric and latent components of LOCOFLOW impose modeling assumptions rather than reveal true lineage relationships. The perturbations are in silico predictions; moreover, the geometric-patch and random-minibatch configurations differ in coupling and training seed. Matched ablations, lineage-resolved measurements, and experimental perturbations are needed to establish which response patterns reflect biological mechanisms. Within these limits, LOCOFLOW provides a framework for studying how local cell population structure shapes inferred trajectories while representing uncertainty in cell fate.

## Reproducibility Statement

The Methods and Appendices A–D describe the transport objectives, bridge construction, time conversions, hierarchical latent variables, and evaluation conventions. Dataset-specific evaluation protocols are given in the experimental section and Appendix D. The supplementary tables report the evaluated ablation configurations and their results; the accompanying text distinguishes training-seed variation from repeated simulation draws and evaluation subsamples.

## Ethics Statement

Computational trajectories and expression perturbations describe model predictions, not clinically validated interventions. Expression-defined neighborhoods do not imply physical cell–cell interactions. Data access and reuse remain subject to the conditions of the original datasets.

## AI Use Statement

We used OpenAI Codex to assist with literature retrieval, research ideation and execution, and drafting and polishing manuscript text. Research assistance included methodological discussion, experimental planning, code implementation (including preprocessing and evaluation), and result interpretation; writing assistance included figure and table formatting. The authors manually verified all AI-assisted outputs incorporated into this work. Scientific claims and conclusions reflect the authors’ independent judgment, and the authors take full responsibility for the accuracy and integrity of the manuscript, code, and reported results.

## A Hierarchical TRANSPORT AND OVERLAP ACCOUNTING

Within a patch pair (*p, q*), the cell-level objective is

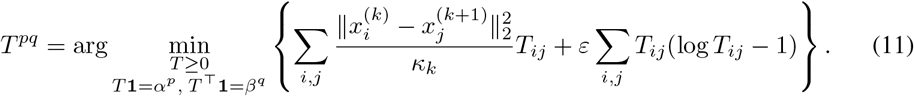

The entropy coefficient is *ε*; the local marginals are defined in Section 3.1. Each local plan is extended by zero outside its patch supports. If patch and cell plans satisfy their prescribed marginals, Eq. (3) gives

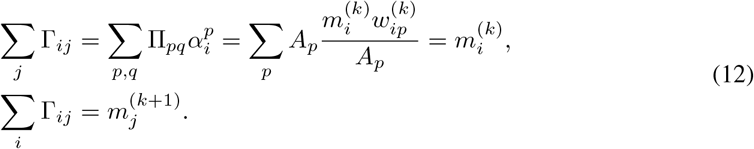

Thus patch overlap does not duplicate mass. Averaging feasible plans with common marginals also preserves these identities. Multiple nonzero destinations in *Q* encode inferred alternatives, not measured ancestry. The source-sampling distribution remains important: uniform source sampling produces target mixture 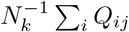, which need not equal the mass-weighted transport marginal 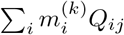

For positive-semidefinite covariances, the squared Bures distance is

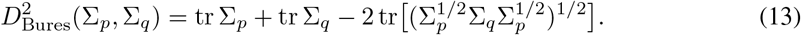

If 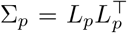 and 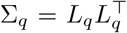, the last trace equals 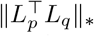, allowing evaluation through low-rank factors. The term compares local spread in addition to the center displacement used by the patch transport cost.

## B Bridge GEOMETRY AND TIME PARAMETERIZATION

Let *e*_ij_ denote fixed nearest-neighbor edge weights in the source patch, and define 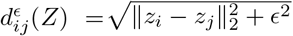, where *ϵ >* 0 stabilizes near-coincident points. The radial penalty is

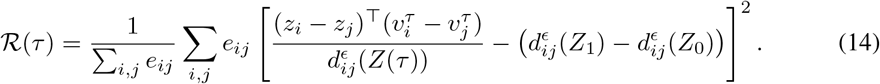

The first term in brackets is exactly 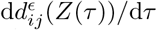. Both distances and relative velocities are invariant to common translation. The endpoint distance difference specifies a prior on deformation per unit local phase; it is not a measurement of intermediate biological speed.

The residual prefactor *τ* (1 − *τ*) in Eq. (4) fixes endpoint positions but not endpoint derivatives. Physical time *t*, interval phase *τ*, and normalized global time *u* are related by

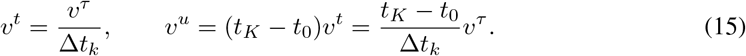

Using the appropriate factor is necessary when intervals have different durations. The geometric penalties restrict admissible paths without guaranteeing exact geodesics, confinement to a known intermediate manifold, or the absence of every trajectory reversal.

## C Target-code INITIALIZATION AND HIERARCHICAL UNCERTAINTY

During initialization, the bridge is fixed and 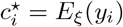 is learned jointly with the velocity network using

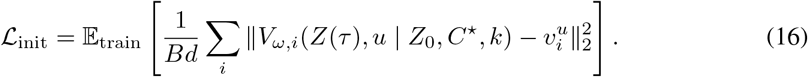

The expectation includes training interval, endpoint, and time sampling with their associated weights. Gradients propagate through the conditioning codes to *E*_*ξ*_. The encoder is frozen after initialization; the velocity parameters initialize hierarchical latent training. Target codes are standardized within each interval using training observations. A patch-level summary contains their mean and standard deviation, while each cell-level posterior receives its corresponding standardized code.

The hierarchical posterior in Eq. (7) and independent Gaussian priors yield

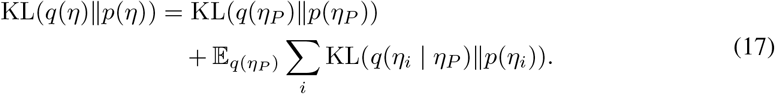

Conditioning inputs are omitted only in this identity. The expectation over the shared latent accounts for dependence of each cell posterior on *η*_*P*_. Neither latent is itself an identified fate label. KL regularization reduces a train–inference distribution mismatch but does not guarantee calibration or rule out posterior collapse. At inference, randomness enters through interval-level latent draws, not through a continuous diffusion increment. Propagating predicted states between intervals is an inference procedure, not an additional rollout loss in Eq. (8).

## D Evaluation CONVENTIONS

Evaluation was performed in the fixed representation and data split specified for each experiment. Model-specific normalization was reversed before scoring: EB and iPSC were evaluated in their standardized benchmark coordinates, *C. elegans* in the released 50-dimensional PCA space, and Axolotl regeneration in normalized spatial coordinates and ten expression PCs.

Distributional agreement was measured using *W*_1_, *W*_2_, and MMD for EB, iPSC, and *C. elegans*. Wasserstein distances were computed by exact optimal transport between uniformly weighted empirical distributions, using Euclidean costs for *W*_1_ and the square root of the optimal squared-Euclidean transport cost for *W*_2_. MMD was reported as

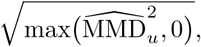

using the Gaussian kernel *k*(*x, y*) = exp[−∥*x − y*∥^2^*/*(2*b*)], where *b* was the lower median of positive pooled pairwise squared distances. For EB and iPSC, scores were averaged over eight LoCoFlow draws per evaluation window and then over windows within each training seed. For *C. elegans*, all 222 founder cells and all observed target cells were used; LoCoFlow scores were averaged over 200 draws and then over the five held-out time points. Deterministic baselines contributed one prediction per source cell.

LARRY fate prediction was evaluated on 859 Day-2 founders against their observed descendant fate distributions using accuracy, balanced accuracy, the predicted undifferentiated fraction (U), and raw Brier score. Accuracy excluded U from the predicted class candidates, with all-U predictions counted as incorrect; balanced accuracy averaged recall over supported ground-truth classes. Raw Brier scores retained all predicted probabilities, including U, with its target probability set to zero. LoCoFlow, VRFM and scDiffEq used 2,000 draws per founder.

Axolotl regeneration evaluation used five fixed 1,024-cell subsamples per seed and reported *W*_2_, untruncated unbiased MMD^2^ averaged over six Gaussian kernels, and the energy-distance V-statistic. For LoCoFlow, 100 endpoints were averaged per source cell before scoring.

Unless explicitly identified as a single-seed analysis, results are means ± sample standard deviations across training seeds: five seeds. Draws and subsamples were not treated as independent training replicates. LARRY ablations and perturbations were reported separately, and perturbation *z* = 0 denoted an intervention rather than an untreated control.

## E LARRY ABLATION RESULTS

A1 sets the KL coefficient to zero; A3 removes hierarchical latent variables; A6 uses random-minibatch OT. These interventions do not separately identify every subcomponent. Both ablation tables report means and sample standard deviations across five training seeds. For each seed, fate probabilities are estimated from 2,000 sampled trajectories per starting cell; the standard deviations summarize variation across training seeds, not across individual trajectories. In particular, A6’s lower Brier coexists with lower balanced accuracy, and should not be interpreted as evidence either for or against geometric context under all criteria. The A1–A6 labels identify the interventions listed in Tables 5 and 6.

**Table 5:**
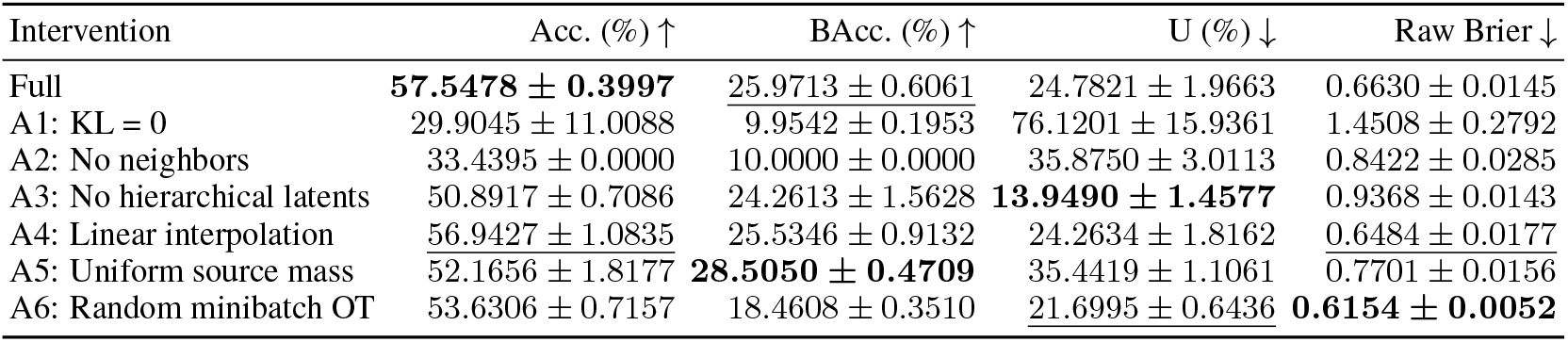
LARRY development geometry-set ablations. Values are means ± sample SD across five training seeds, with 2,000 trajectories per starting cell per seed. Bold and underlining mark the best and second-best means, respectively. Lower U is better in this evaluation.

**Table 6:**
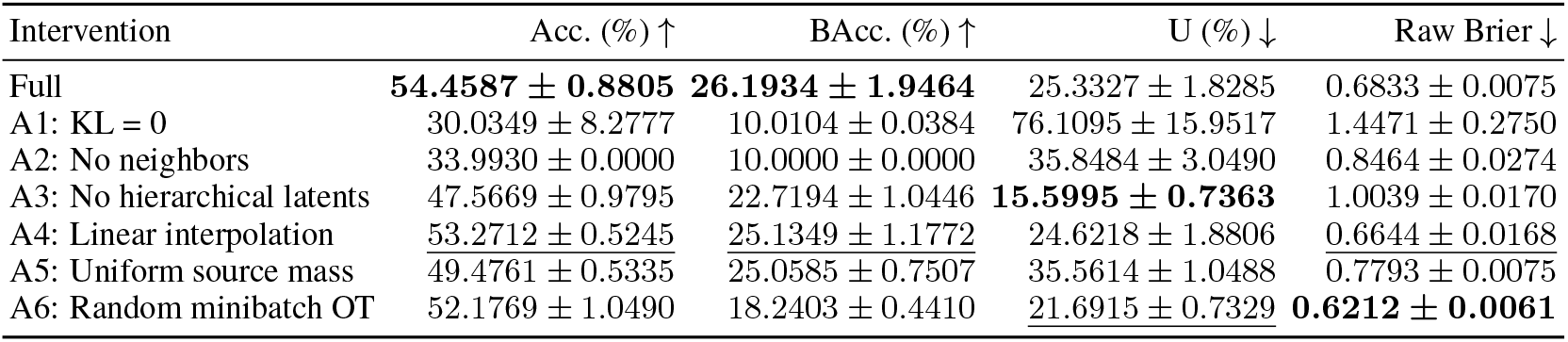
LARRY test-set ablations. Values are means ± sample SD across five training seeds, with 2,000 trajectories per starting cell per seed. Bold and underlining mark the best and second-best means, respectively. Lower U is better in this evaluation.

## F Supplementary FIGURES

The supplementary figures provide population views, representative lineage examples, dose responses, a single-gene screen, and spatial predictions. Figure 2 and the *z* = 0 intervention in Figure 3 appear in the main text. Other dose-specific panels are shown here. Observed clones are untreated references, not ground-truth responses to the computational interventions. Projection axes describe state-space embeddings rather than developmental time or measured lineage paths.

**Figure S01:**
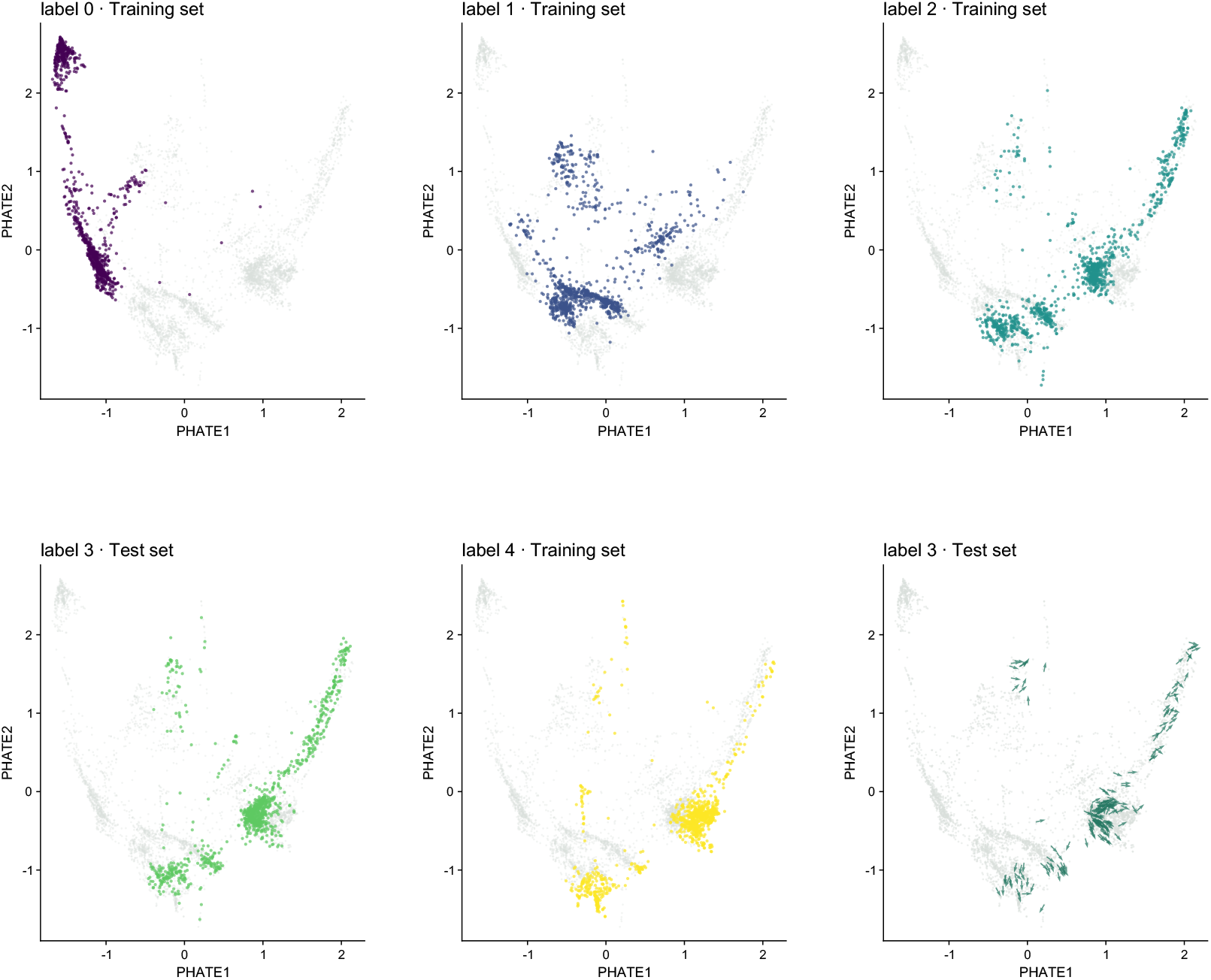
Observed EB states and the supplied reference-velocity view. Panels show labels 0, 1, 2, and 4 as training observations and label 3 as test observations in PHATE coordinates. The final panel overlays reference arrows at label 3; these are not asserted to be directly measured cellular velocities.

**Figure S02:**
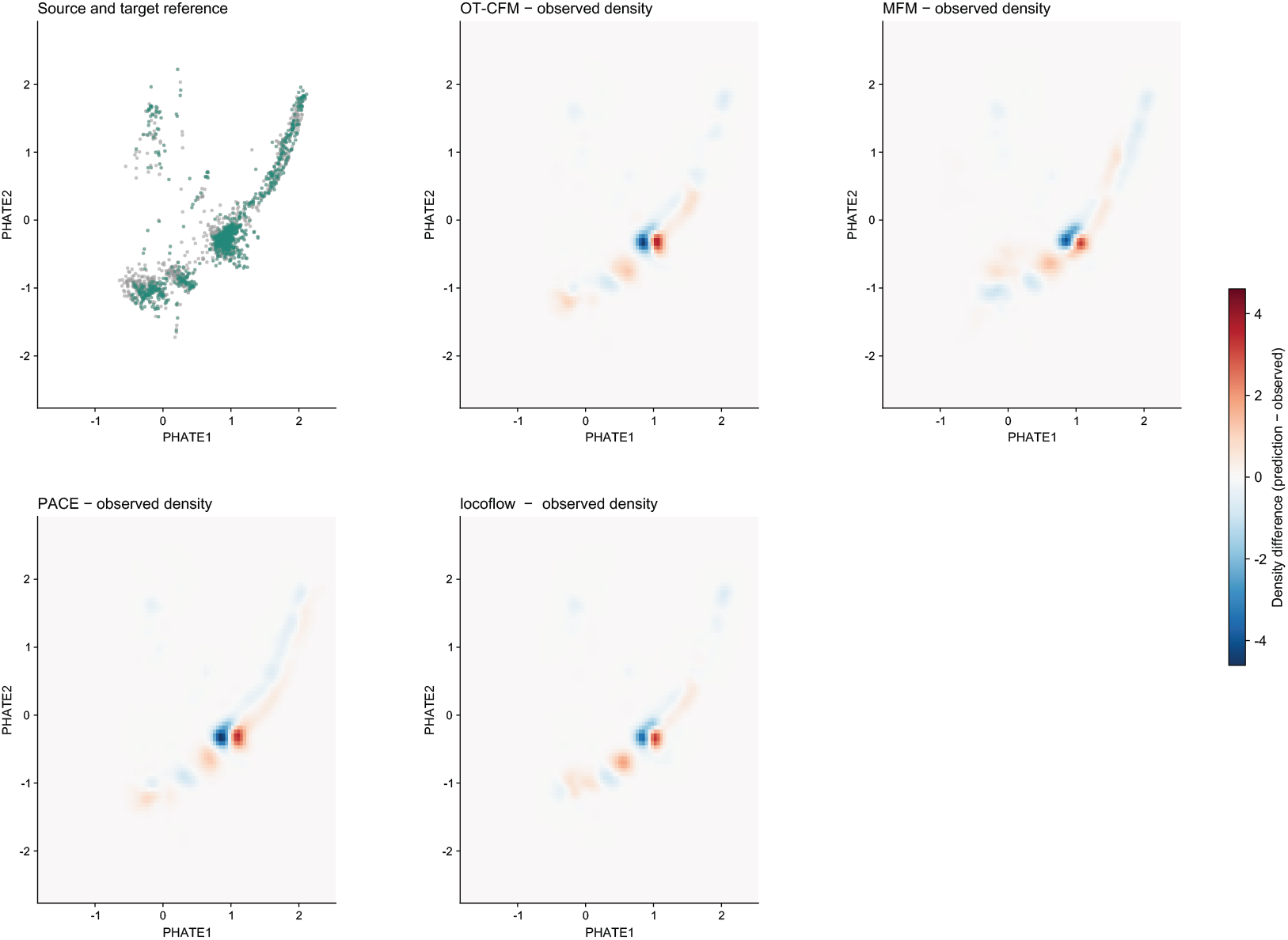
EB density differences. The reference panel is followed by predicted-minus-observed density maps for the supplied methods. The diverging color scale identifies excess and deficient density in the displayed coordinates; it is not a high-dimensional error metric.

**Figure S03:**
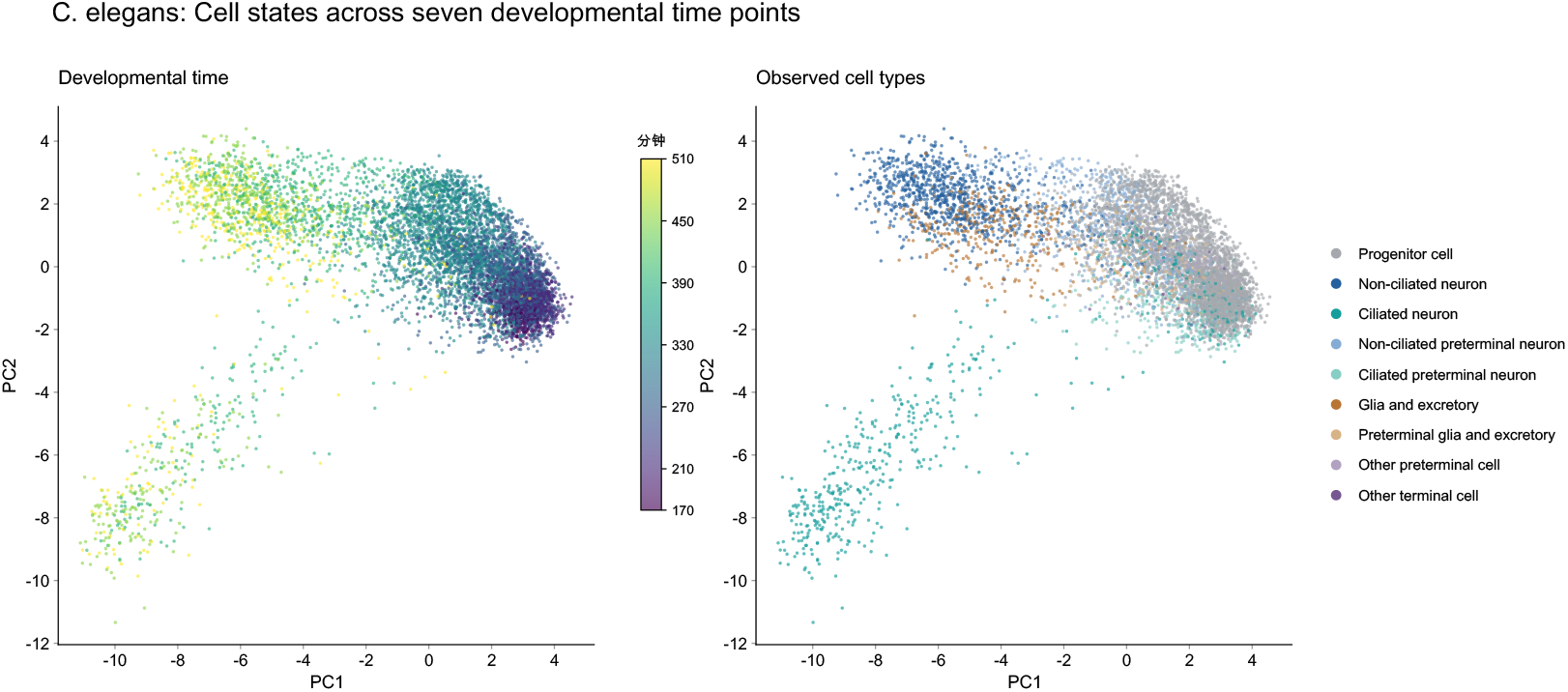
Observed states in the worm dataset. The same *C. elegans* principal-component view is colored by developmental time (minutes; the original color-bar unit is retained) and annotated cell type. This is observed context, not a prediction.

**Figure S04:**
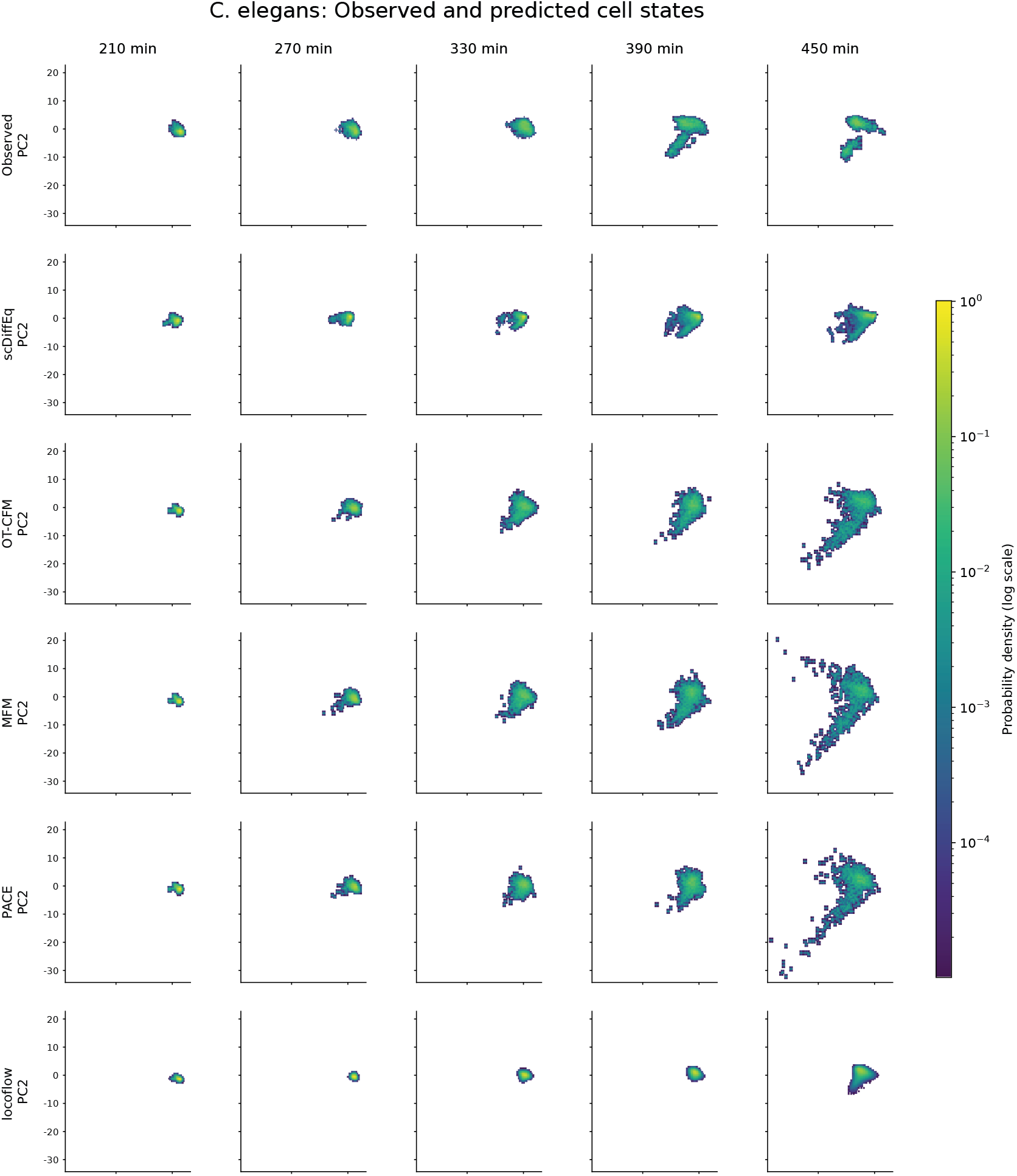
Temporal holdout. In *C. elegans*, columns correspond to 210, 270, 330, 390, and 450 minutes; rows identify observations and model variants. The shared logarithmic color scale represents cell density in PC1/PC2. All method rows and the full color bar are displayed together.

**Figure S06:**
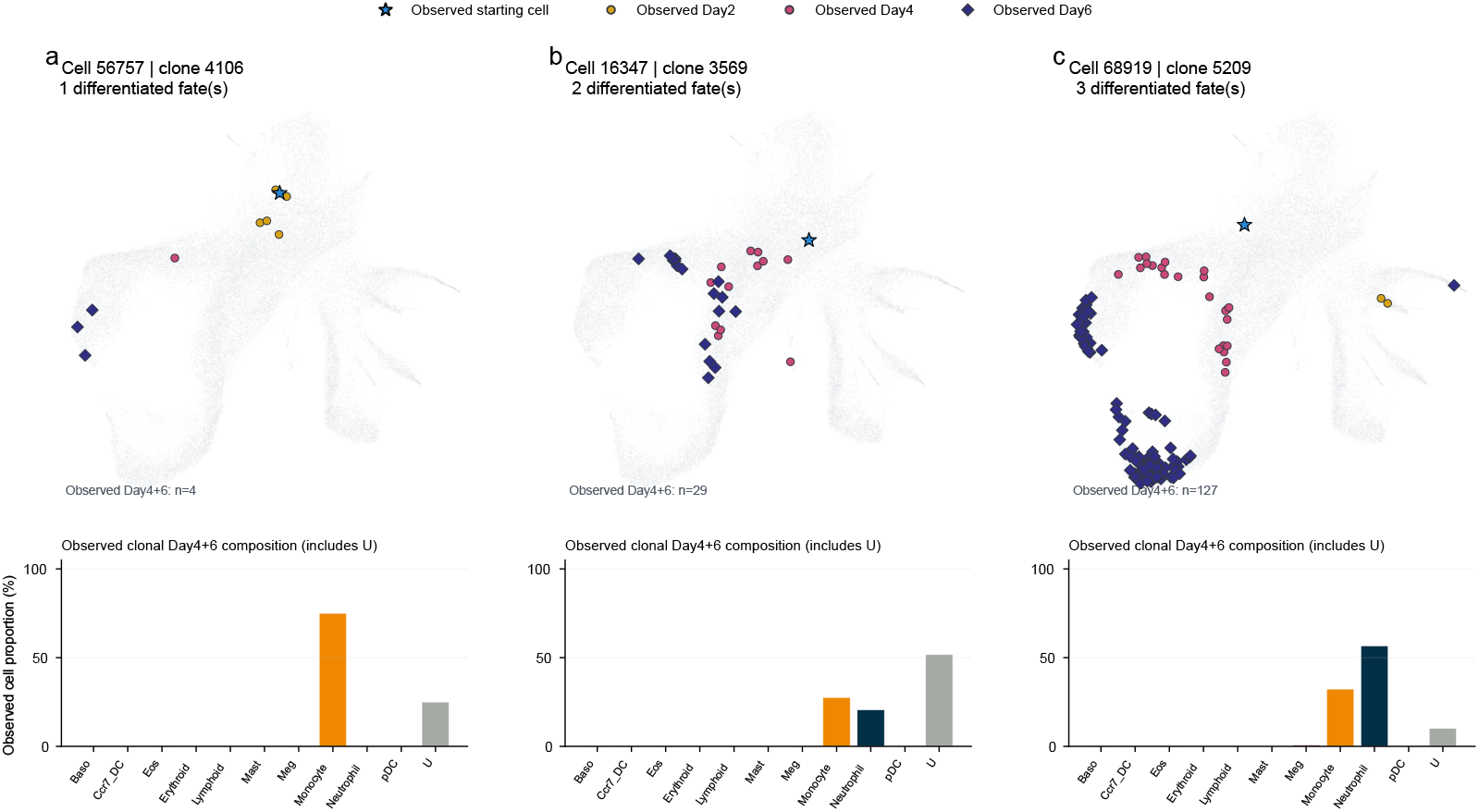
Observed representative LARRY progenitors. Cells 56757 (clone 4106), 16347 (clone 3569), and 68919 (clone 5209) have one, two, and three observed differentiated fates, respectively. Their displayed Day 4+6 descendant counts are 4, 29, and 127. Bars retain U. Selection uses observed fate distributions, not model performance; the internal Figure 2 title is retained from the source.

**Figure S07a:**
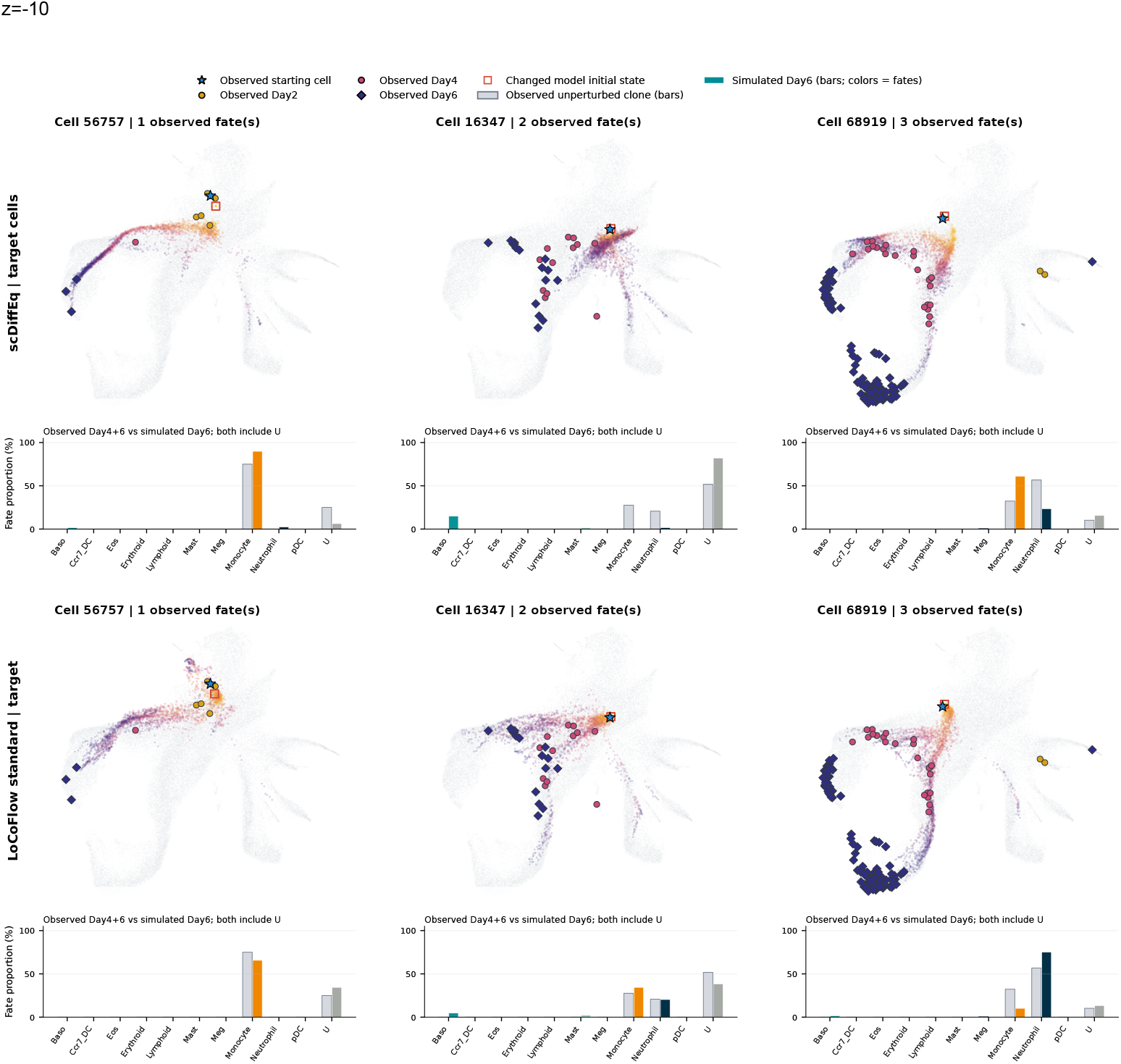
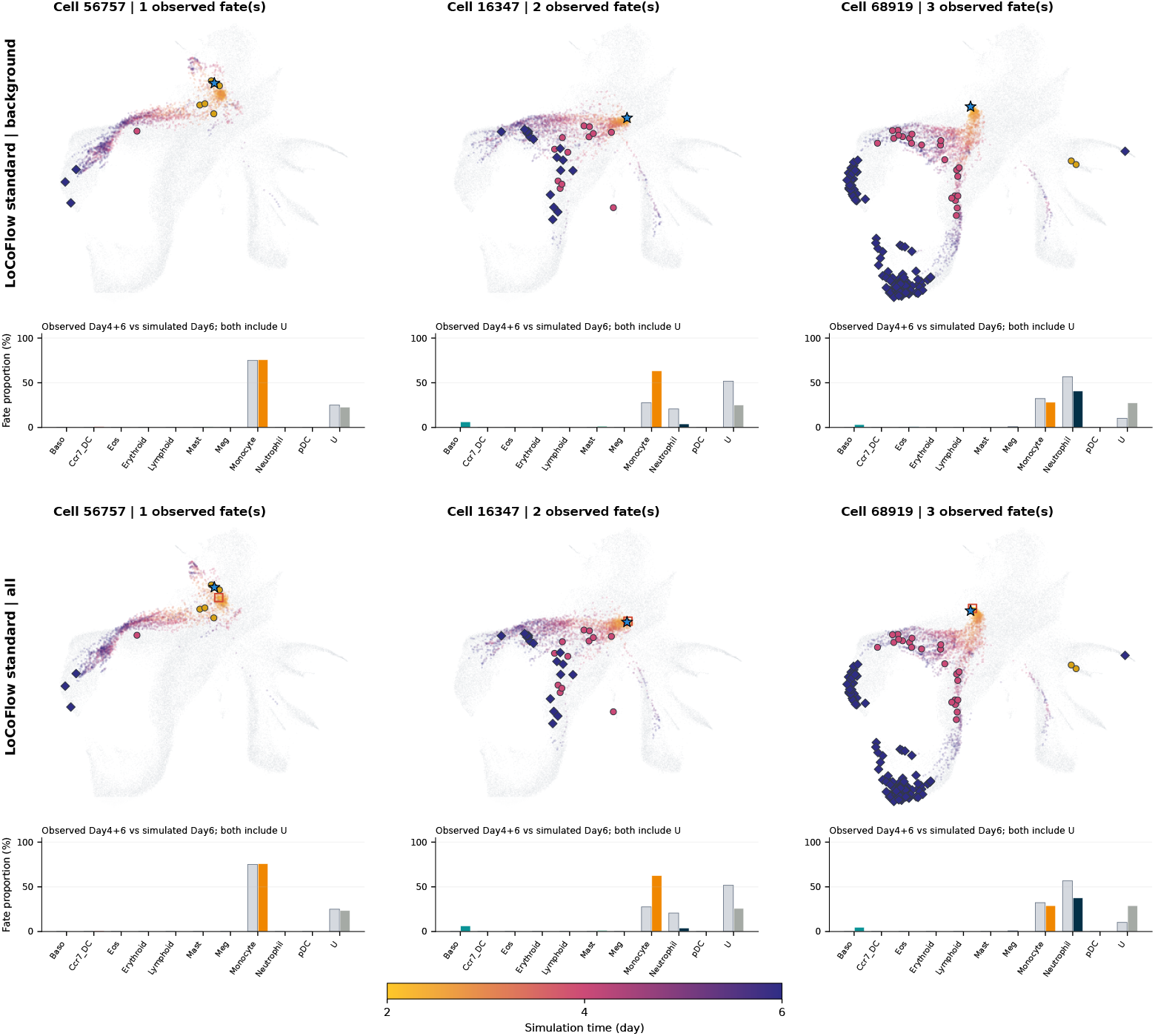
Representative LARRY responses at *z* = −10. Columns show cells 56757, 16347, and 68919 with one, two, and three observed differentiated fates. This part shows scDiffEq target-cell perturbation and LOCOFLOW target-only perturbation; background-only and all-member arms follow. Observed bars pool untreated Day 4+6 descendants, while simulated bars summarize Day 6; both retain U. Points show simulated states, with time encoded by color. The joint clamp affects *Cebpe, Dach1, Lmo4*, and *Mxd1*. These observed clones are not dose-specific ground truth. LOCOFLOW background-only and all-member arms at *z* = −10. Definitions, observed references, and starting-cell columns are unchanged.

**Figure S07b:**
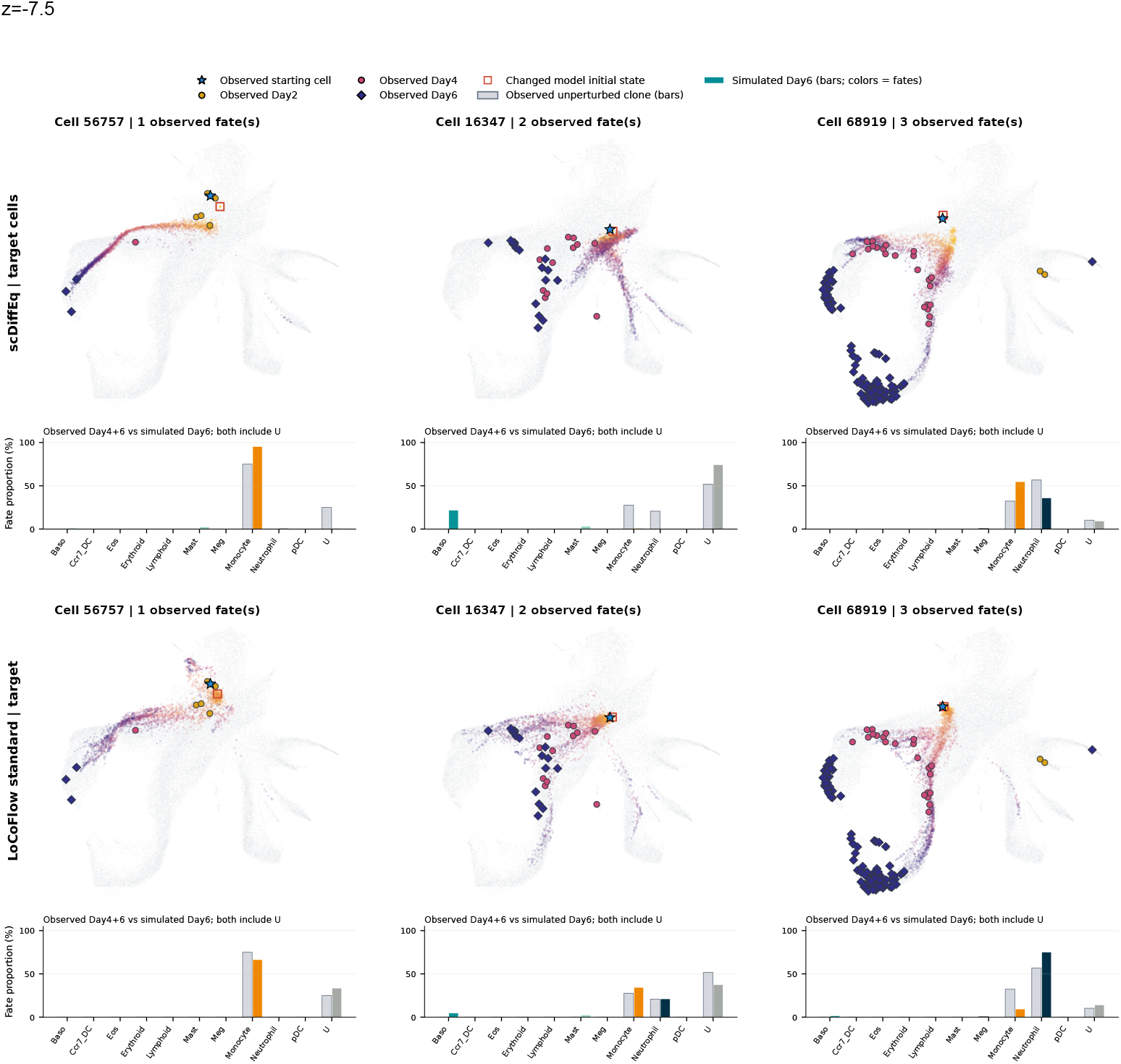
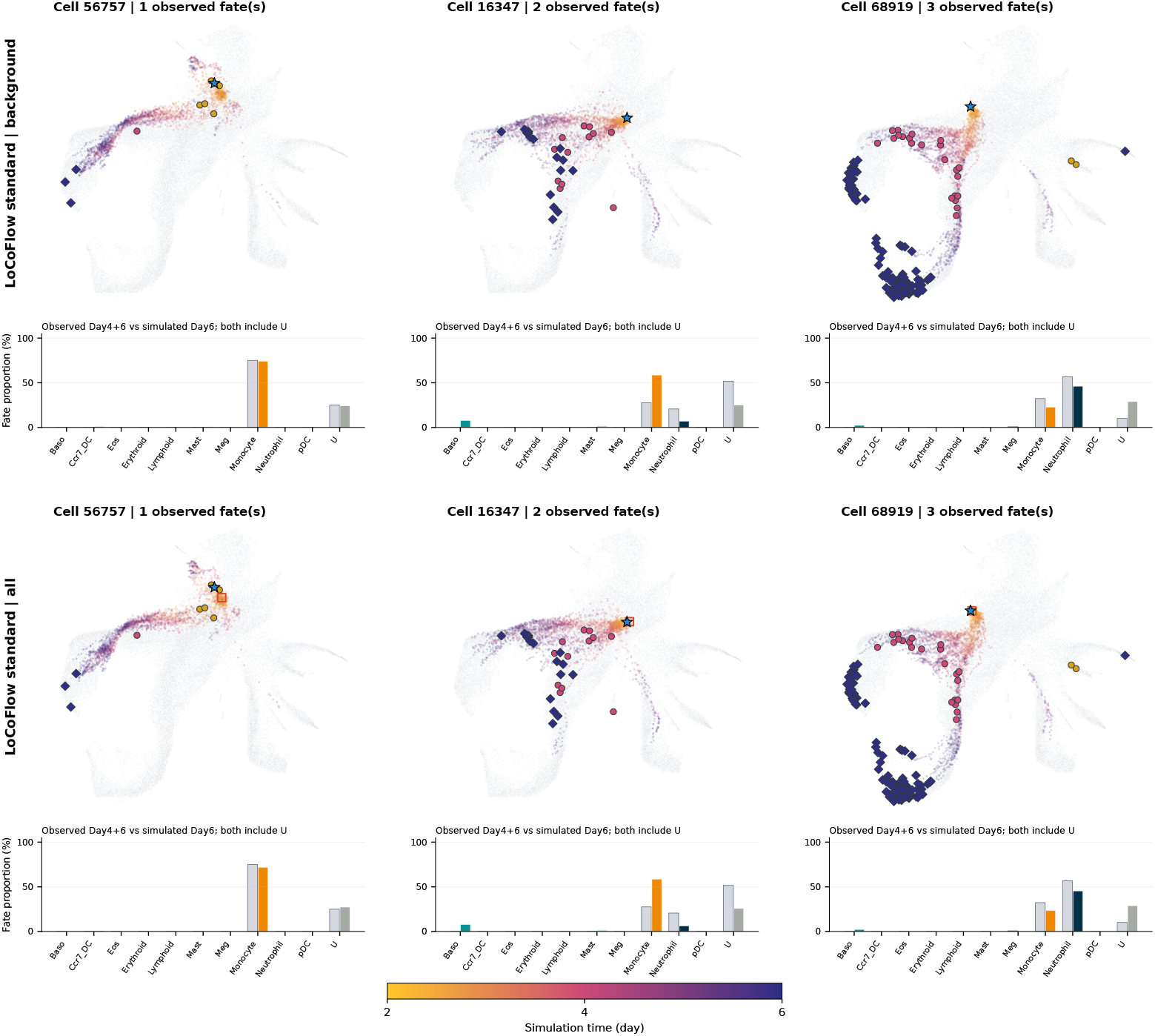
Representative LARRY responses at *z* = −7.5. Columns show cells 56757, 16347, and 68919 with one, two, and three observed differentiated fates. This part shows scDiffEq target-cell perturbation and LOCOFLOW target-only perturbation; background-only and all-member arms follow. Observed bars pool untreated Day 4+6 descendants, while simulated bars summarize Day 6; both retain U. Points show simulated states, with time encoded by color. The joint clamp affects *Cebpe, Dach1, Lmo4*, and *Mxd1*. These observed clones are not dose-specific ground truth. LOCOFLOW background-only and all-member arms at *z* = −7.5. Definitions, observed references, and starting-cell columns are unchanged.

**Figure S07c:**
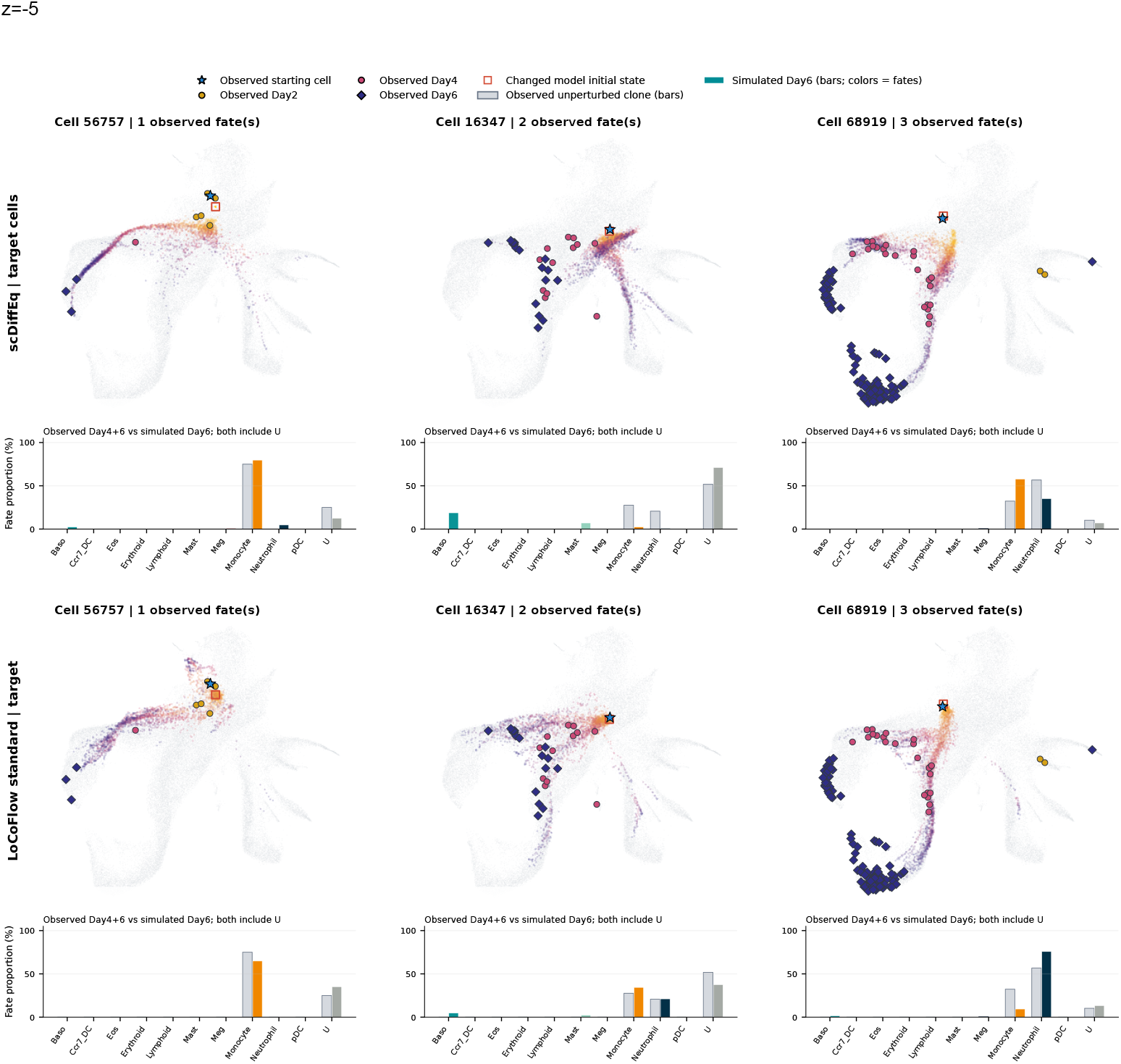
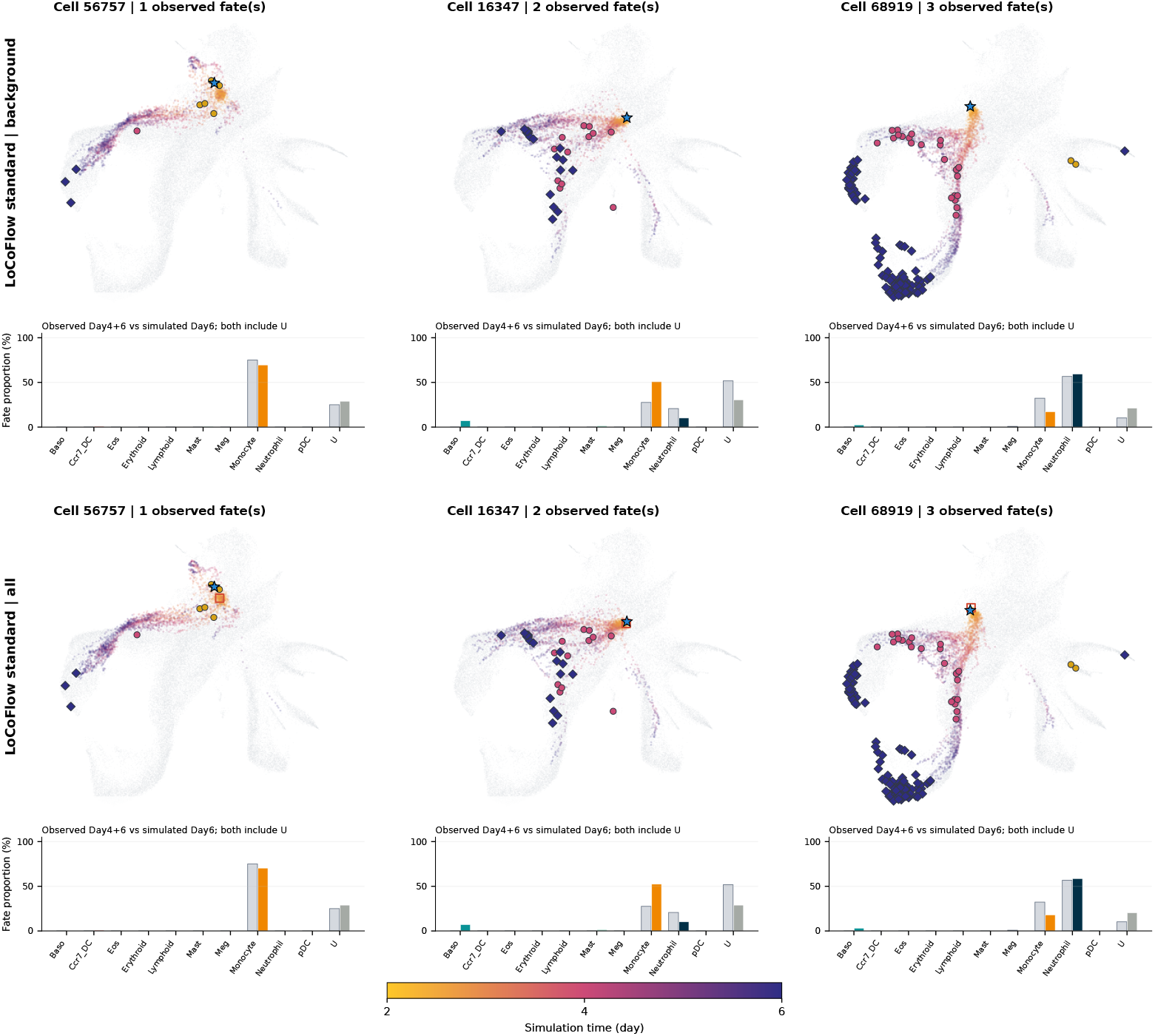
Representative LARRY responses at *z* = −5. Columns show cells 56757, 16347, and 68919 with one, two, and three observed differentiated fates. This part shows scDiffEq target-cell perturbation and LOCOFLOW target-only perturbation; background-only and all-member arms follow. Observed bars pool untreated Day 4+6 descendants, while simulated bars summarize Day 6; both retain U. Points show simulated states, with time encoded by color. The joint clamp affects *Cebpe, Dach1, Lmo4*, and *Mxd1*. These observed clones are not dose-specific ground truth. LOCOFLOW background-only and all-member arms at *z* = −5. Definitions, observed references, and starting-cell columns are unchanged.

**Figure S07d:**
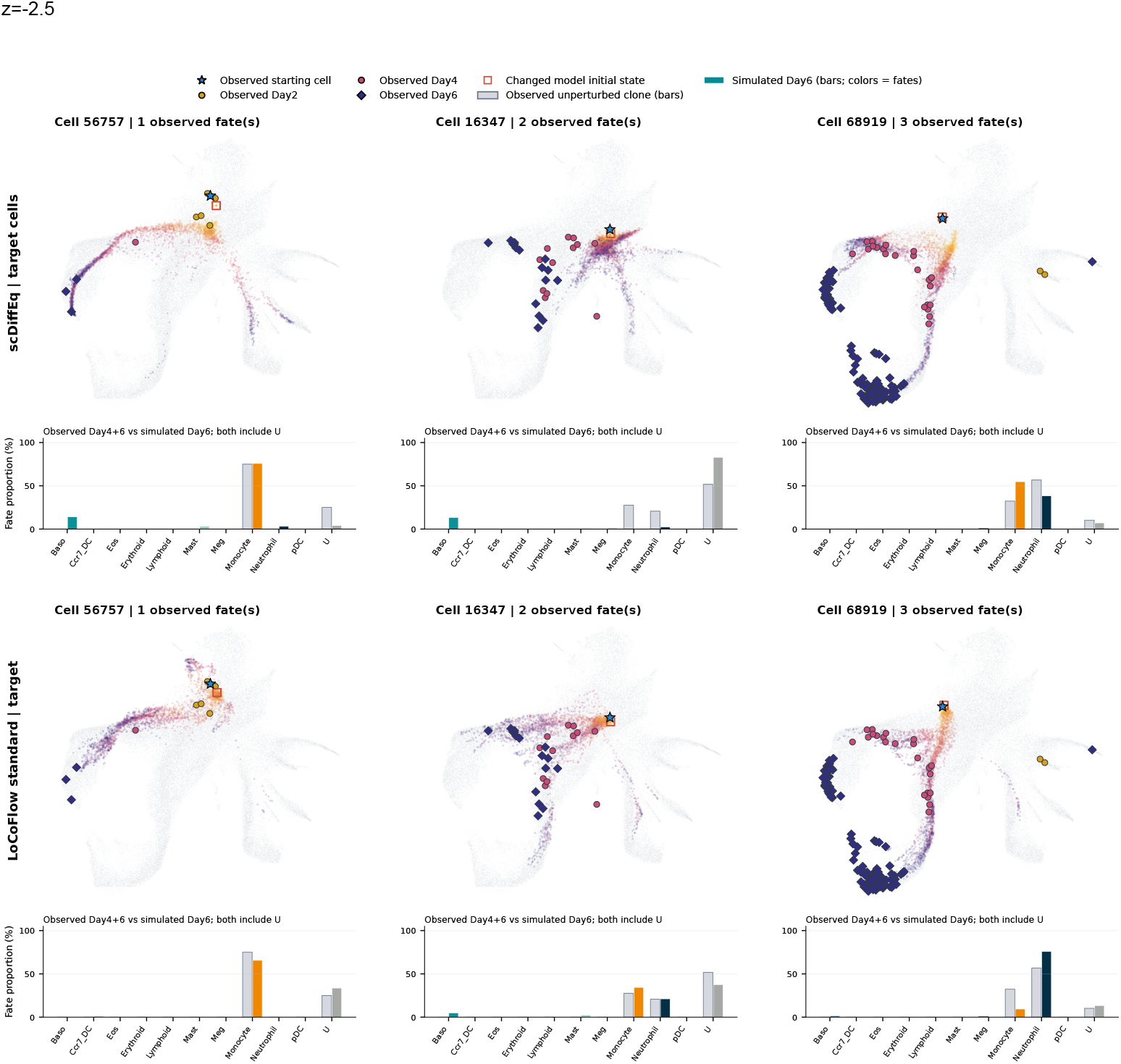
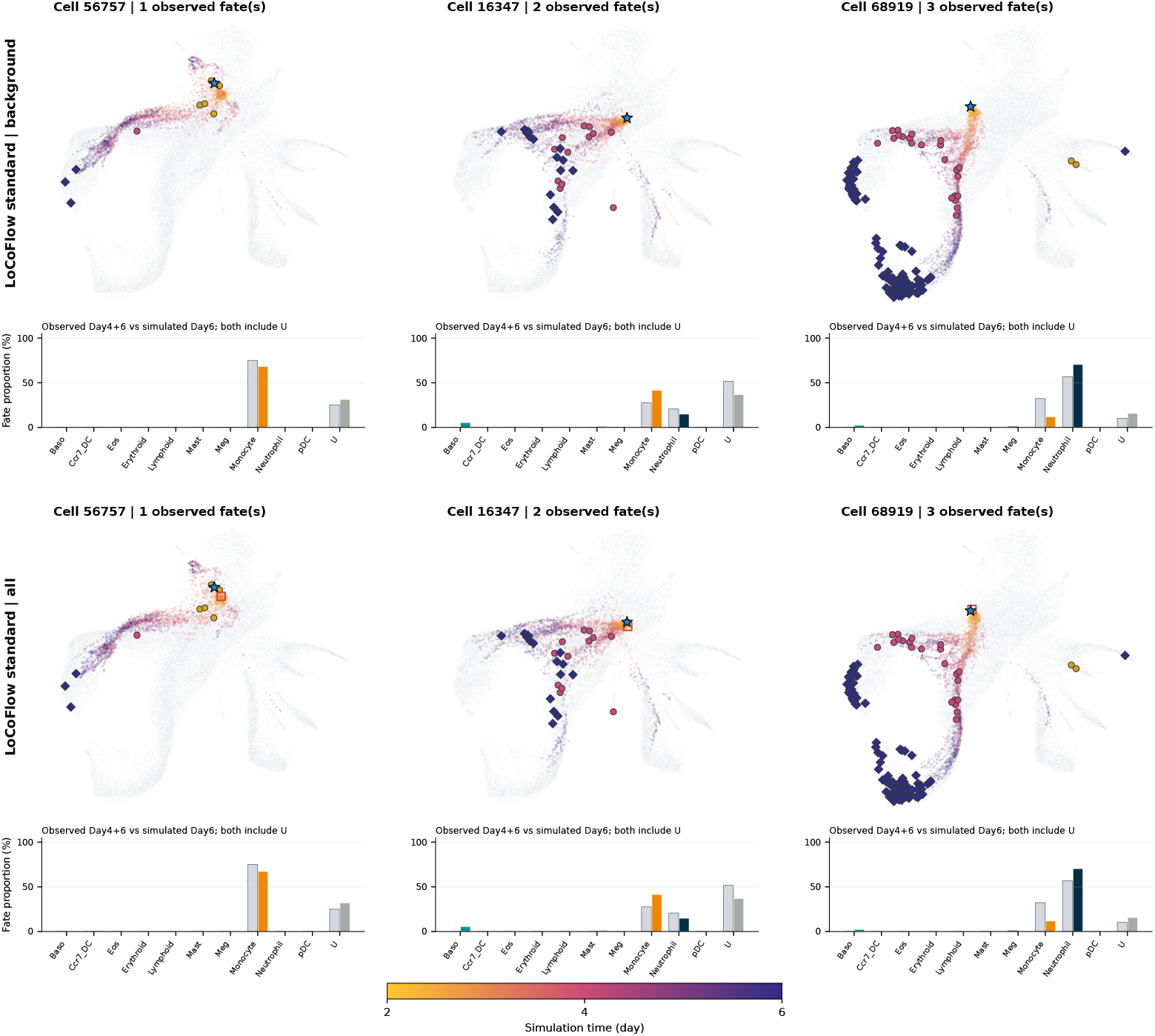
Representative LARRY responses at *z* = −2.5. Columns show cells 56757, 16347, and 68919 with one, two, and three observed differentiated fates. This part shows scDiffEq target-cell perturbation and LOCOFLOW target-only perturbation; background-only and all-member arms follow. Observed bars pool untreated Day 4+6 descendants, while simulated bars summarize Day 6; both retain U. Points show simulated states, with time encoded by color. The joint clamp affects *Cebpe, Dach1, Lmo4*, and *Mxd1*. These observed clones are not dose-specific ground truth. LOCOFLOW background-only and all-member arms at *z* = −2.5. Definitions, observed references, and starting-cell columns are unchanged.

**Figure S07f:**
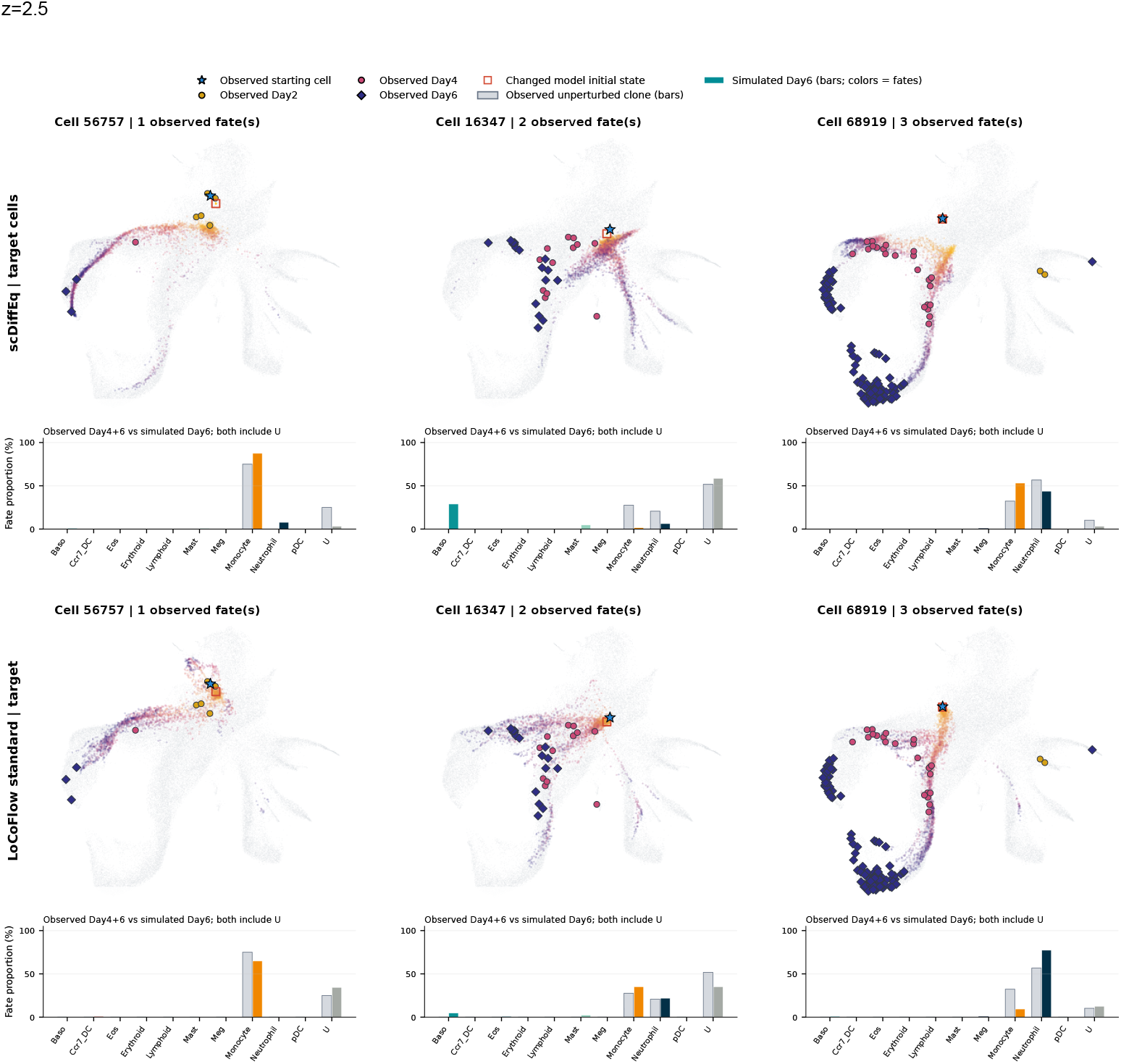
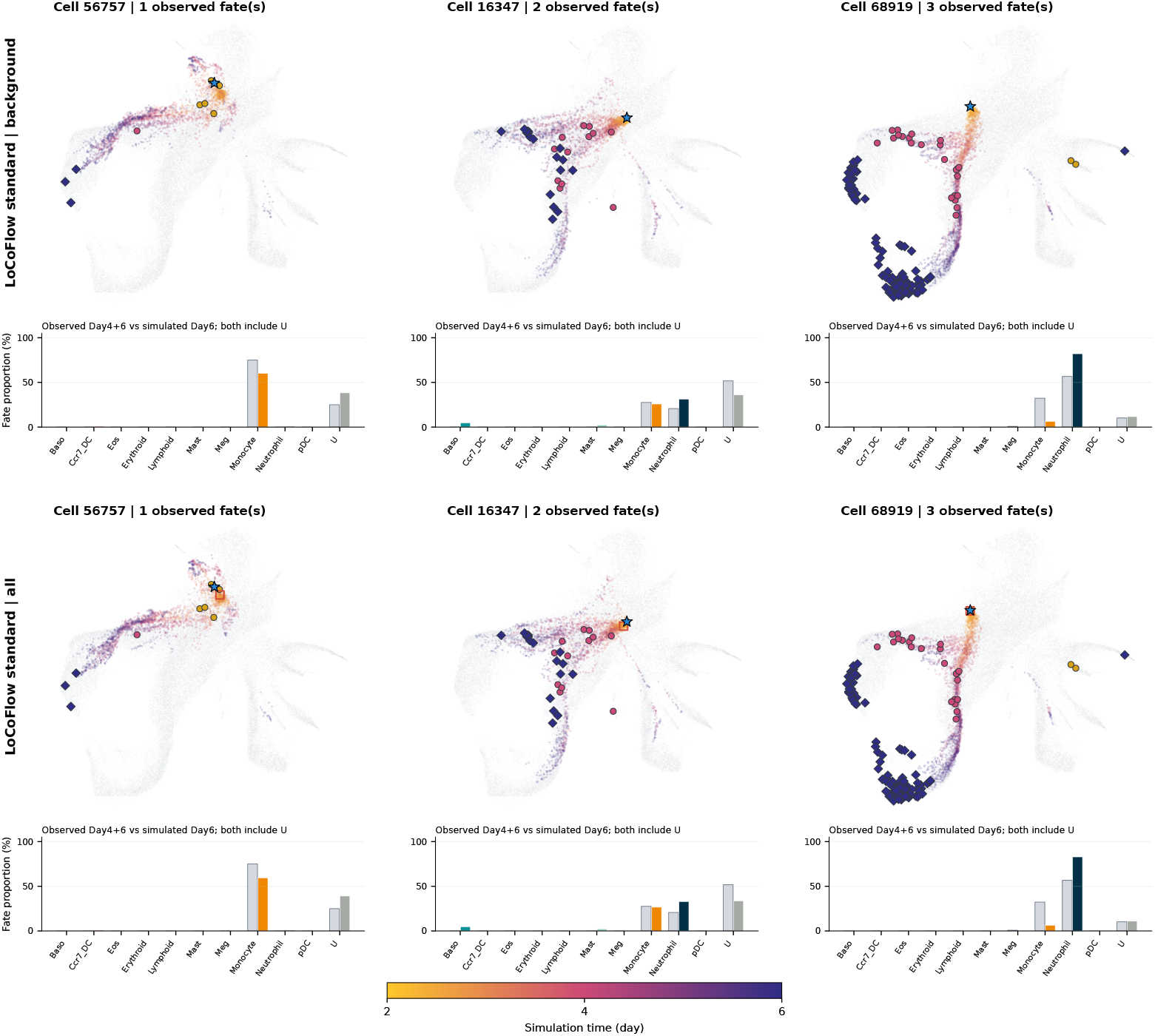
Representative LARRY responses at *z* = 2.5. Columns show cells 56757, 16347, and 68919 with one, two, and three observed differentiated fates. This part shows scDiffEq target-cell perturbation and LOCOFLOW target-only perturbation; background-only and all-member arms follow. Observed bars pool untreated Day 4+6 descendants, while simulated bars summarize Day 6; both retain U. Points show simulated states, with time encoded by color. The joint clamp affects *Cebpe, Dach1, Lmo4*, and *Mxd1*. These observed clones are not dose-specific ground truth. LOCOFLOW background-only and all-member arms at *z* = 2.5. Definitions, observed references, and starting-cell columns are unchanged.

**Figure S07g:**
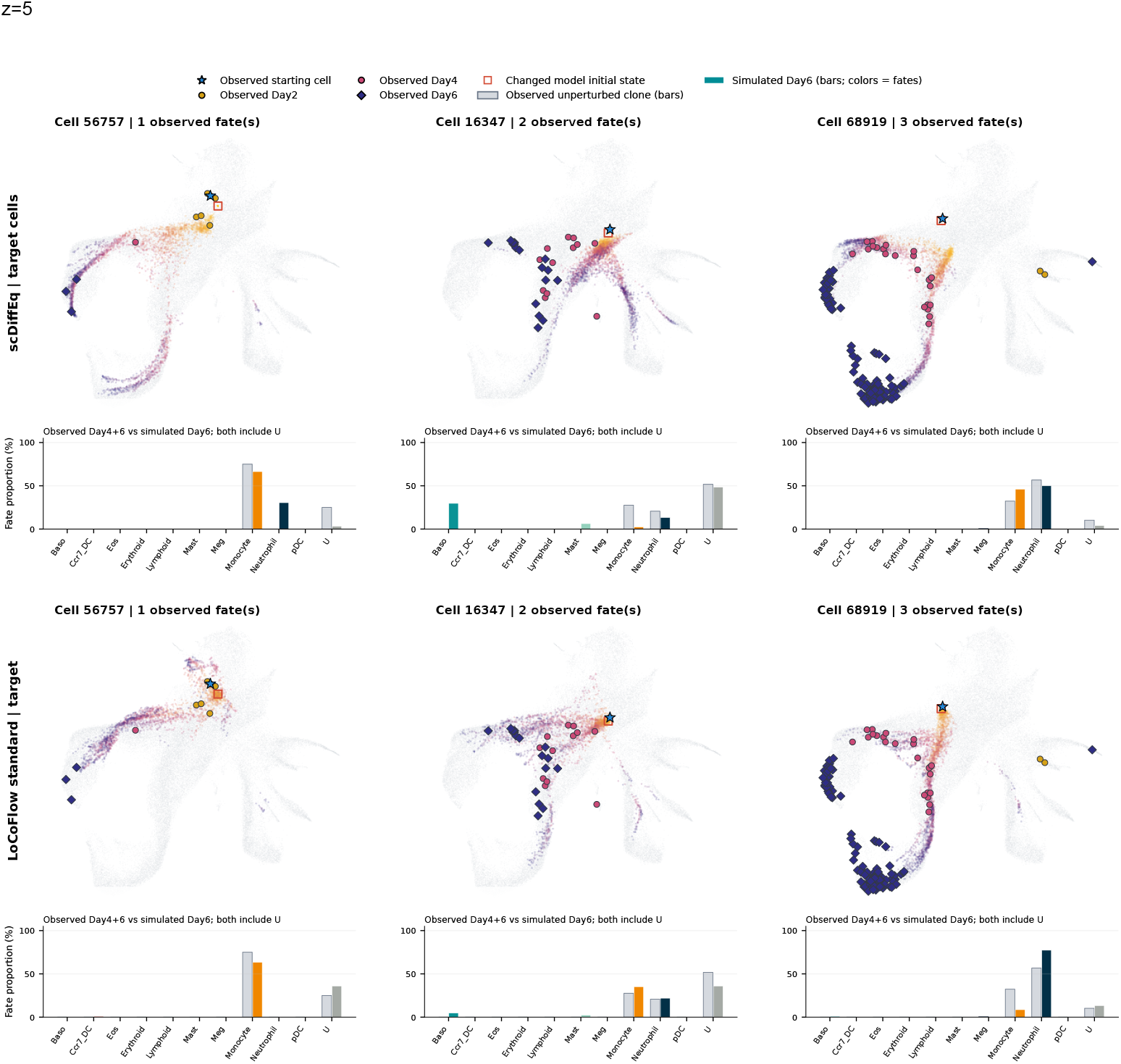
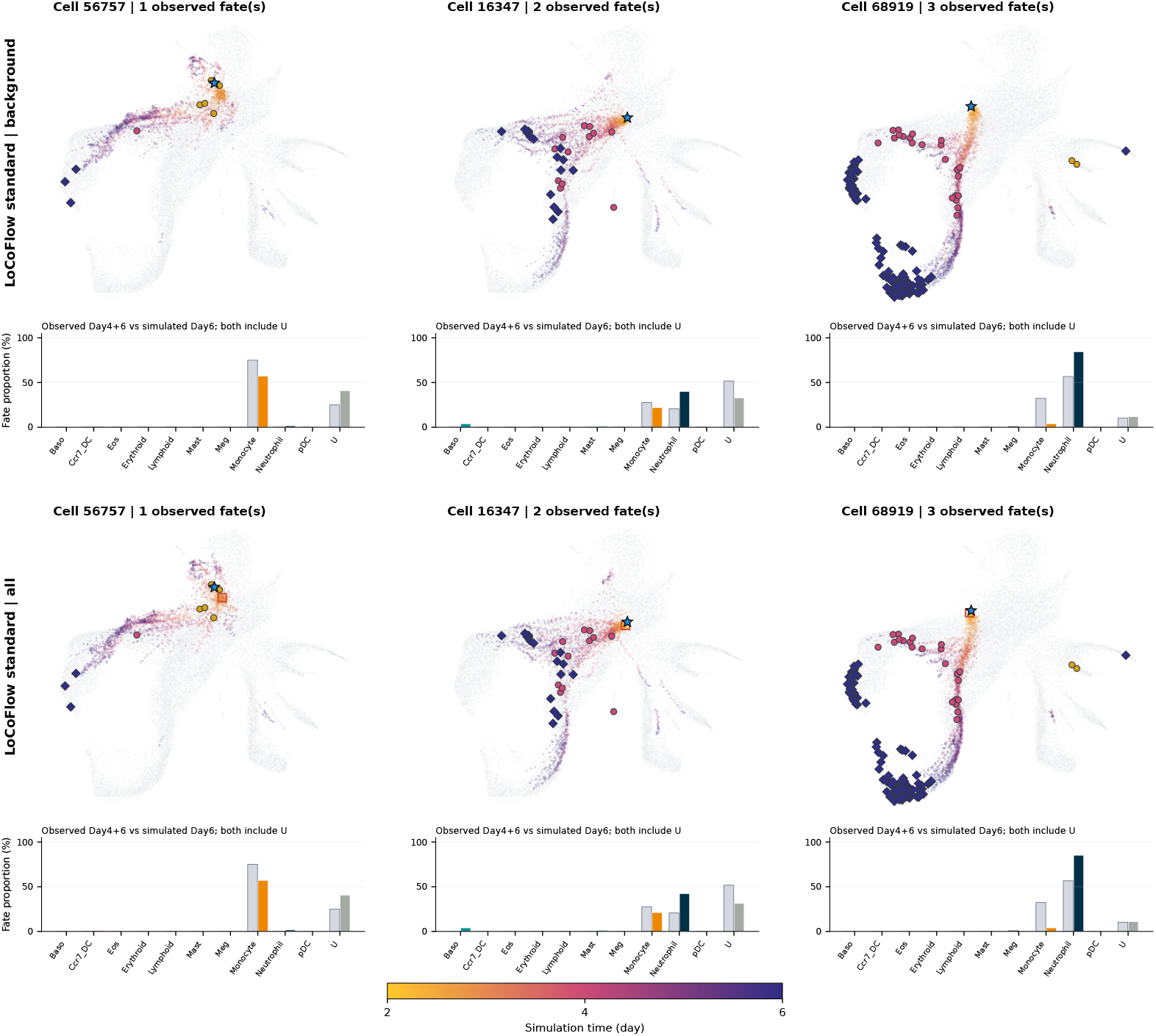
Representative LARRY responses at *z* = 5. Columns show cells 56757, 16347, and 68919 with one, two, and three observed differentiated fates. This part shows scDiffEq target-cell perturbation and LOCOFLOW target-only perturbation; background-only and all-member arms follow. Observed bars pool untreated Day 4+6 descendants, while simulated bars summarize Day 6; both retain U. Points show simulated states, with time encoded by color. The joint clamp affects *Cebpe, Dach1, Lmo4*, and *Mxd1*. These observed clones are not dose-specific ground truth. LOCOFLOW background-only and all-member arms at *z* = 5. Definitions, observed references, and starting-cell columns are unchanged.

**Figure S07h:**
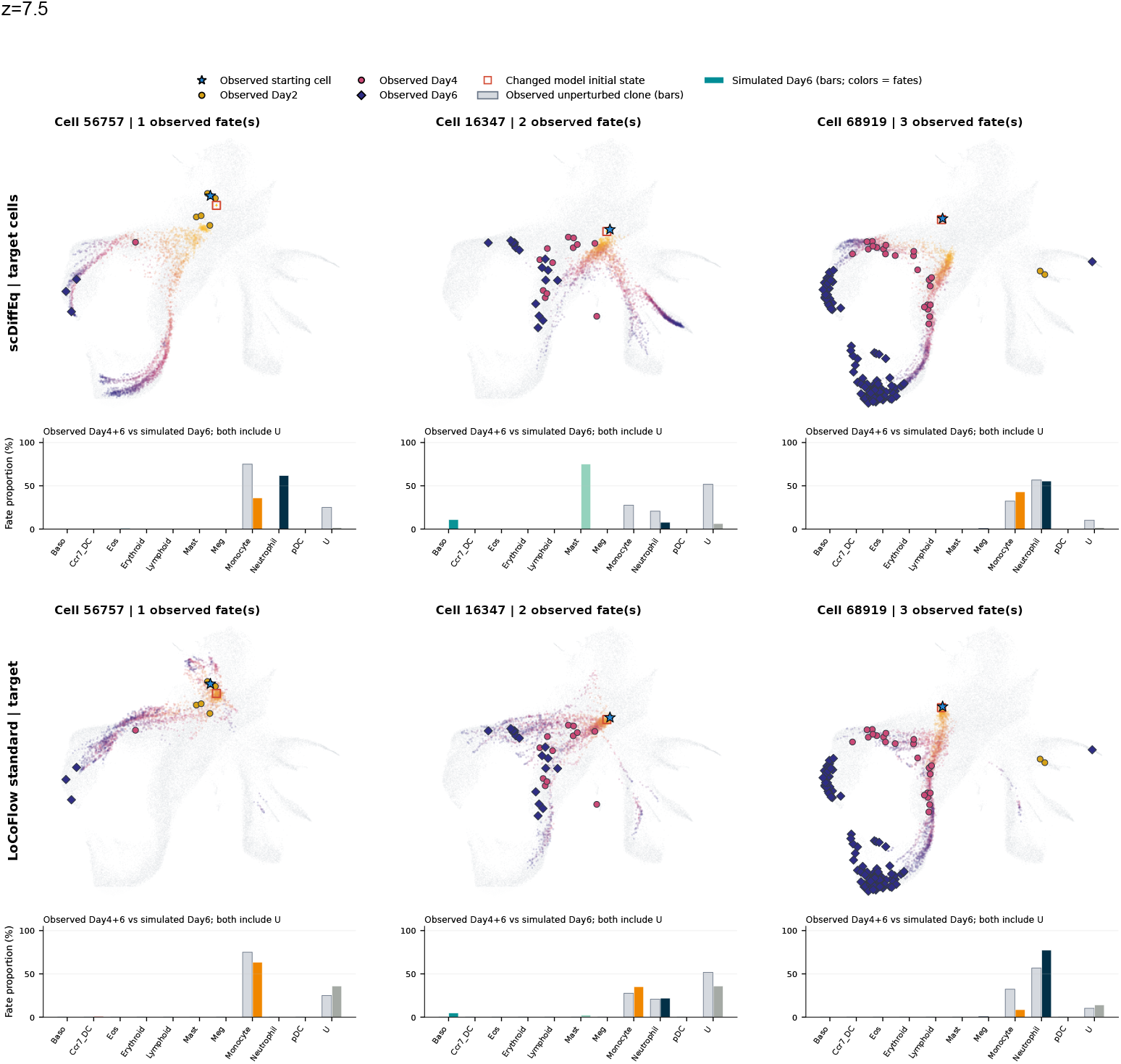
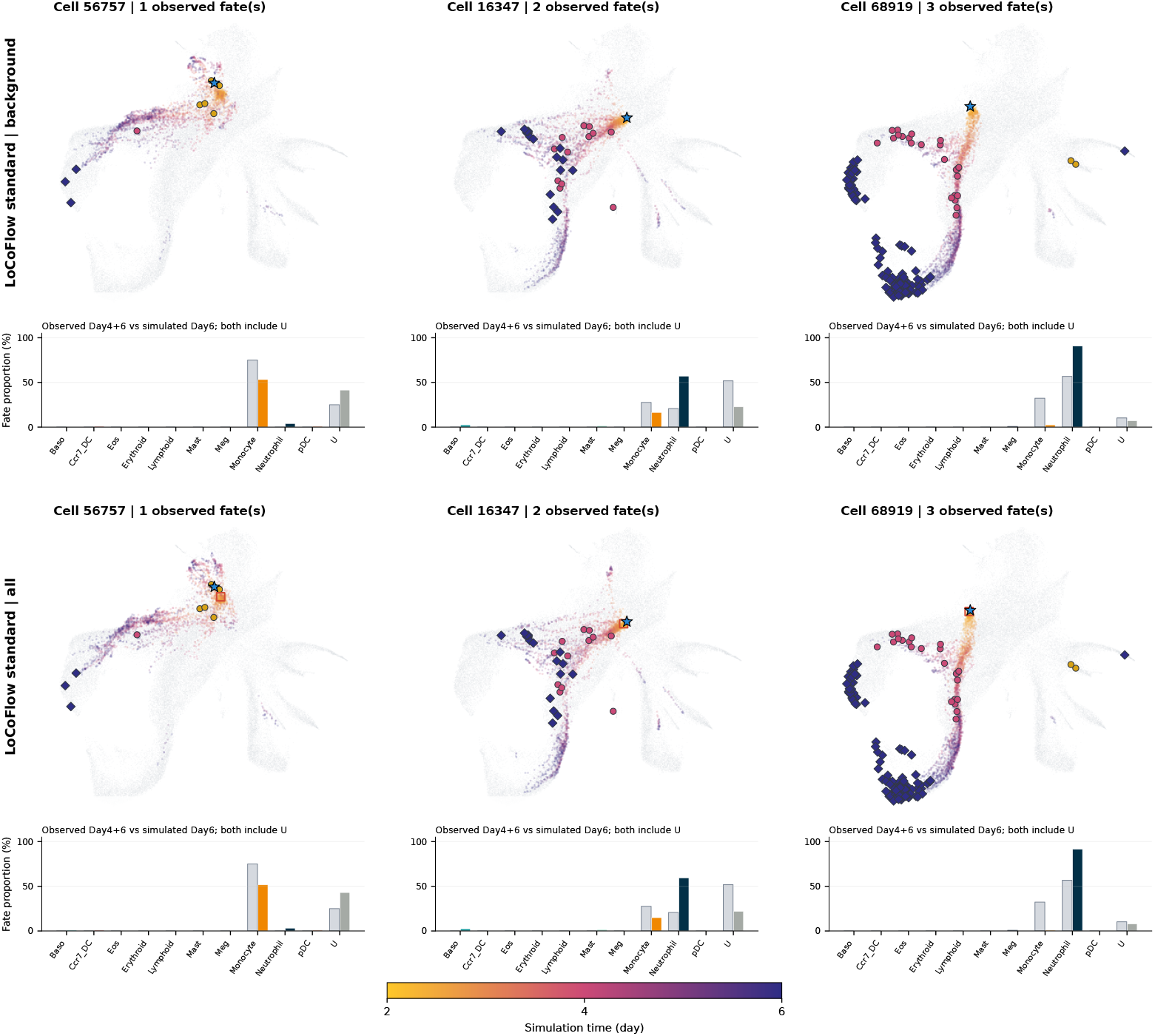
Representative LARRY responses at *z* = 7.5. Columns show cells 56757, 16347, and 68919 with one, two, and three observed differentiated fates. This part shows scDiffEq target-cell perturbation and LOCOFLOW target-only perturbation; background-only and all-member arms follow. Observed bars pool untreated Day 4+6 descendants, while simulated bars summarize Day 6; both retain U. Points show simulated states, with time encoded by color. The joint clamp affects *Cebpe, Dach1, Lmo4*, and *Mxd1*. These observed clones are not dose-specific ground truth. LOCOFLOW background-only and all-member arms at *z* = 7.5. Definitions, observed references, and starting-cell columns are unchanged

**Figure S07i:**
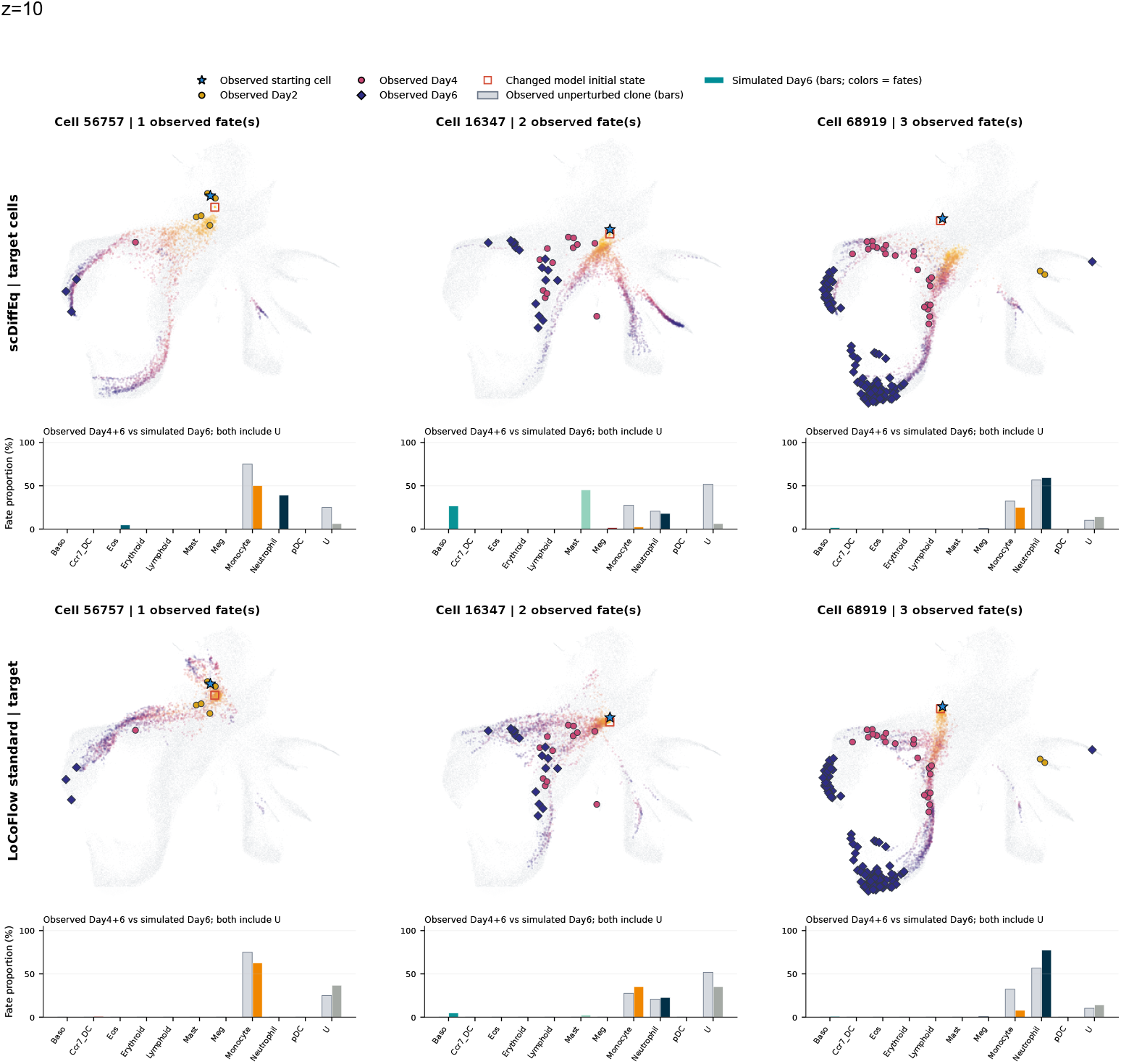
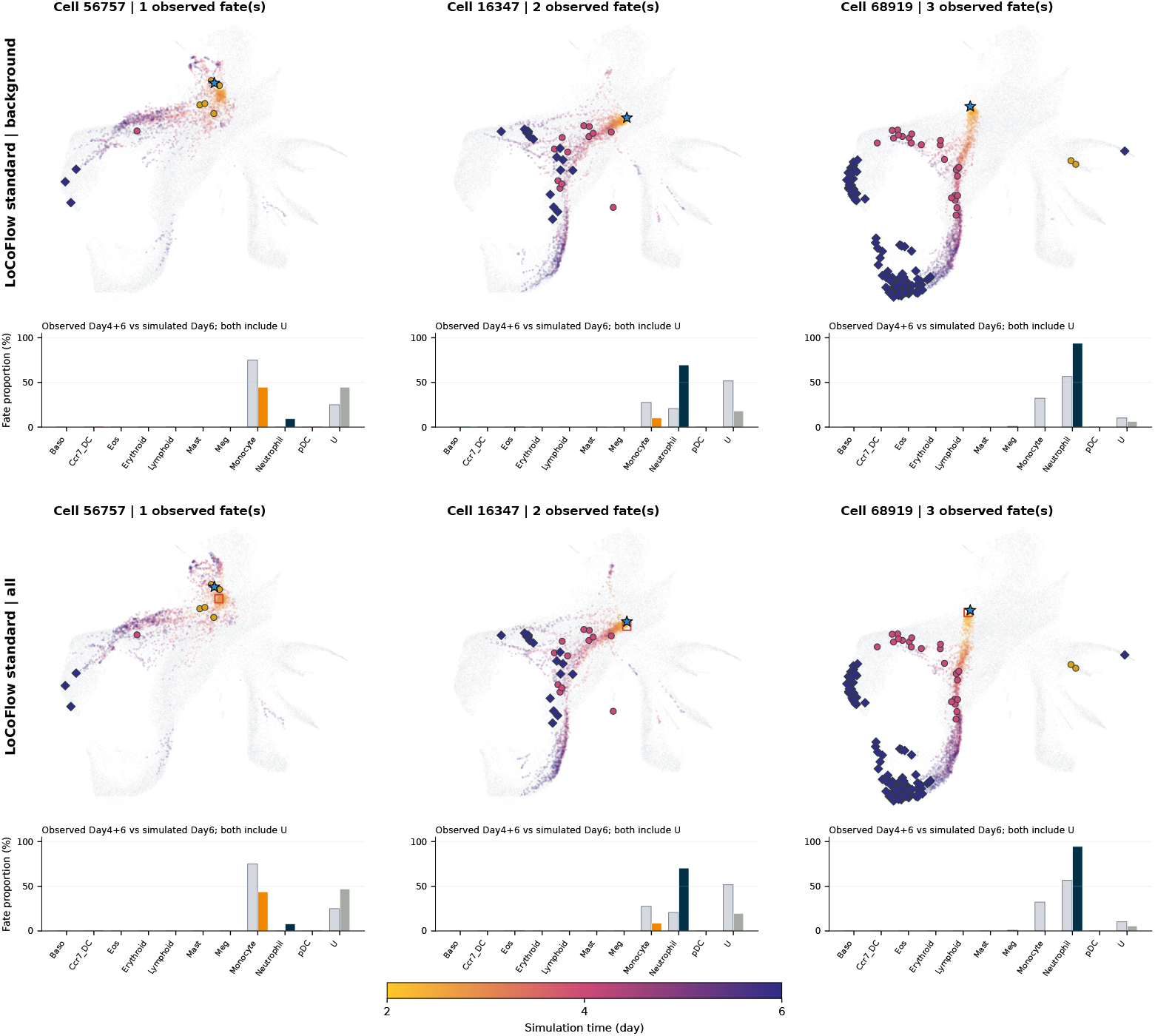
Representative LARRY responses at *z* = 10. Columns show cells 56757, 16347, and 68919 with one, two, and three observed differentiated fates. This part shows scDiffEq target-cell perturbation and LOCOFLOW target-only perturbation; background-only and all-member arms follow. Observed bars pool untreated Day 4+6 descendants, while simulated bars summarize Day 6; both retain U. Points show simulated states, with time encoded by color. The joint clamp affects *Cebpe, Dach1, Lmo4*, and *Mxd1*. These observed clones are not dose-specific ground truth. LOCOFLOW background-only and all-member arms at *z* = 10. Definitions, observed references, and starting-cell columns are unchanged.

**Figure S08:**
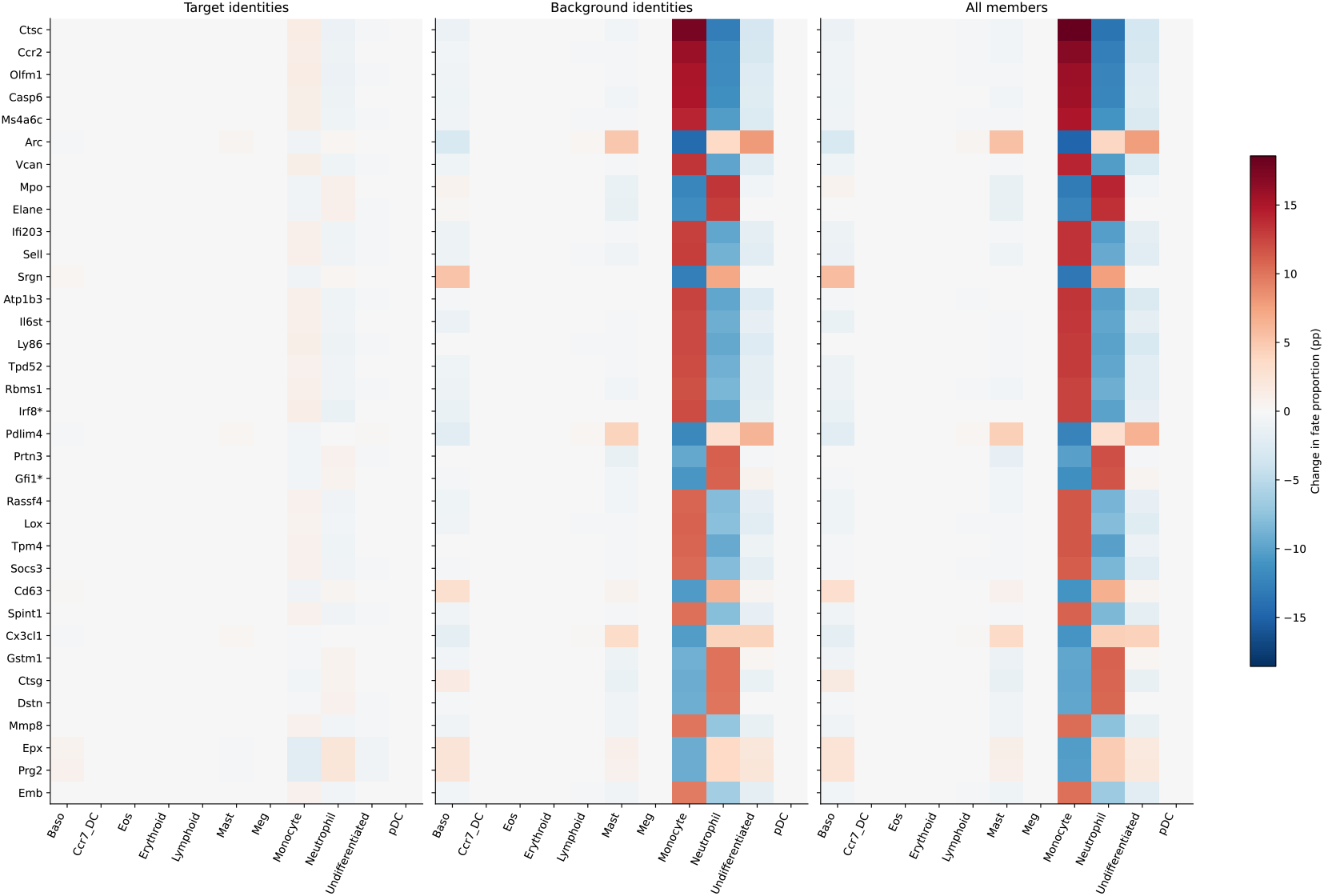
Exploratory single-gene perturbation screen. The supplied heatmap shows 35 selected genes under target-only, background-only, and all-member perturbation. Labels specify standard seed 3, single-gene *z* = 10, and an average over 20 independent noise streams. Asterisks mark membership in the source transcription-factor list. Effect-based gene selection and simulation replication do not establish biological significance or a regulatory mechanism.

**Figure S09:**
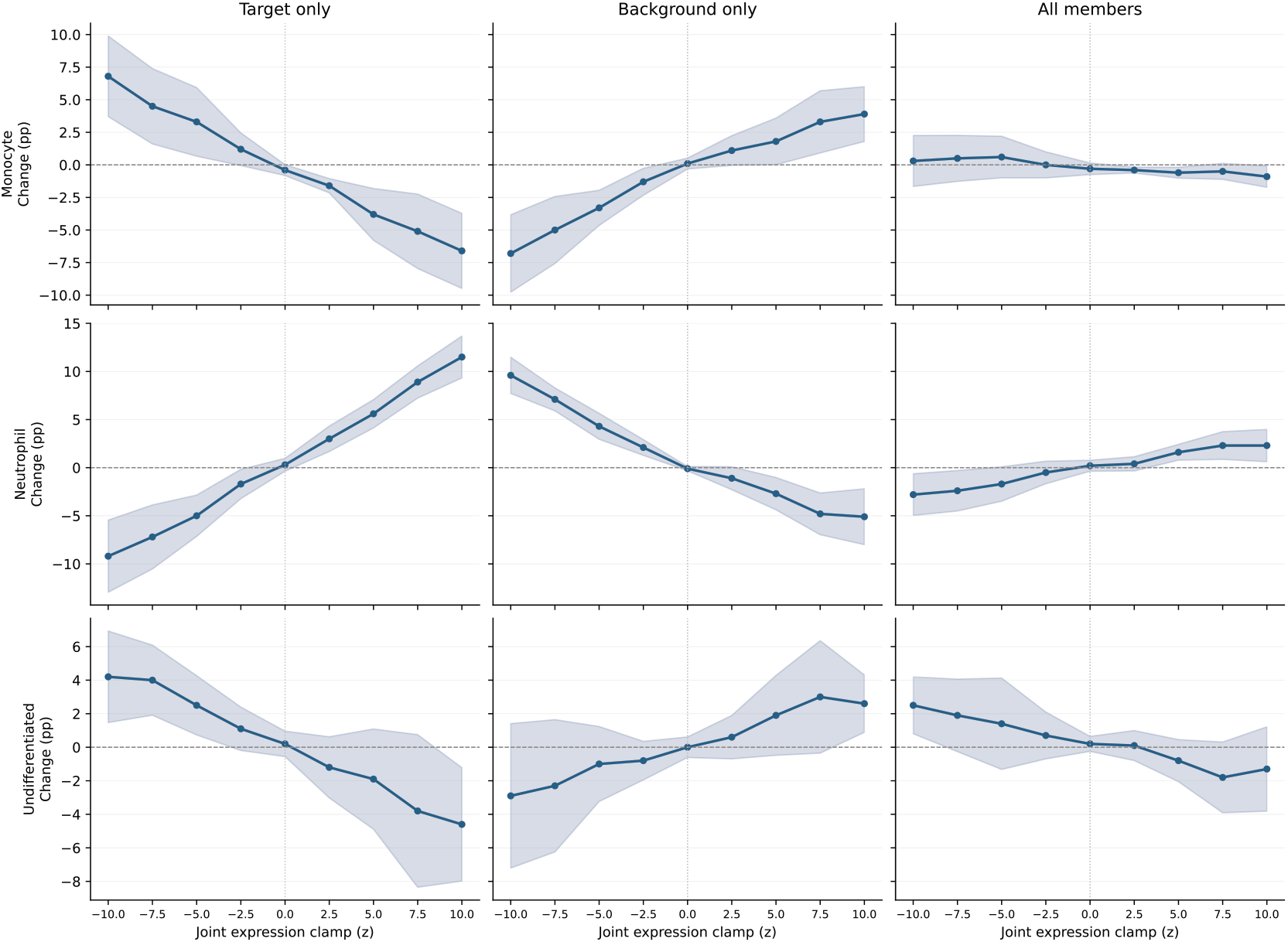
Random-context LARRY responses. Q2 minibatch OT, seed 0, uses one target and 31 random Day 2 backgrounds. Columns are target only, background only, and all members; rows show monocyte, neutrophil, and U changes in percentage points. Target and background effects have opposing directions. Shading is SD across five simulation groups, not training seeds. *z* = 0 remains an intervention.

**Figure S10:**
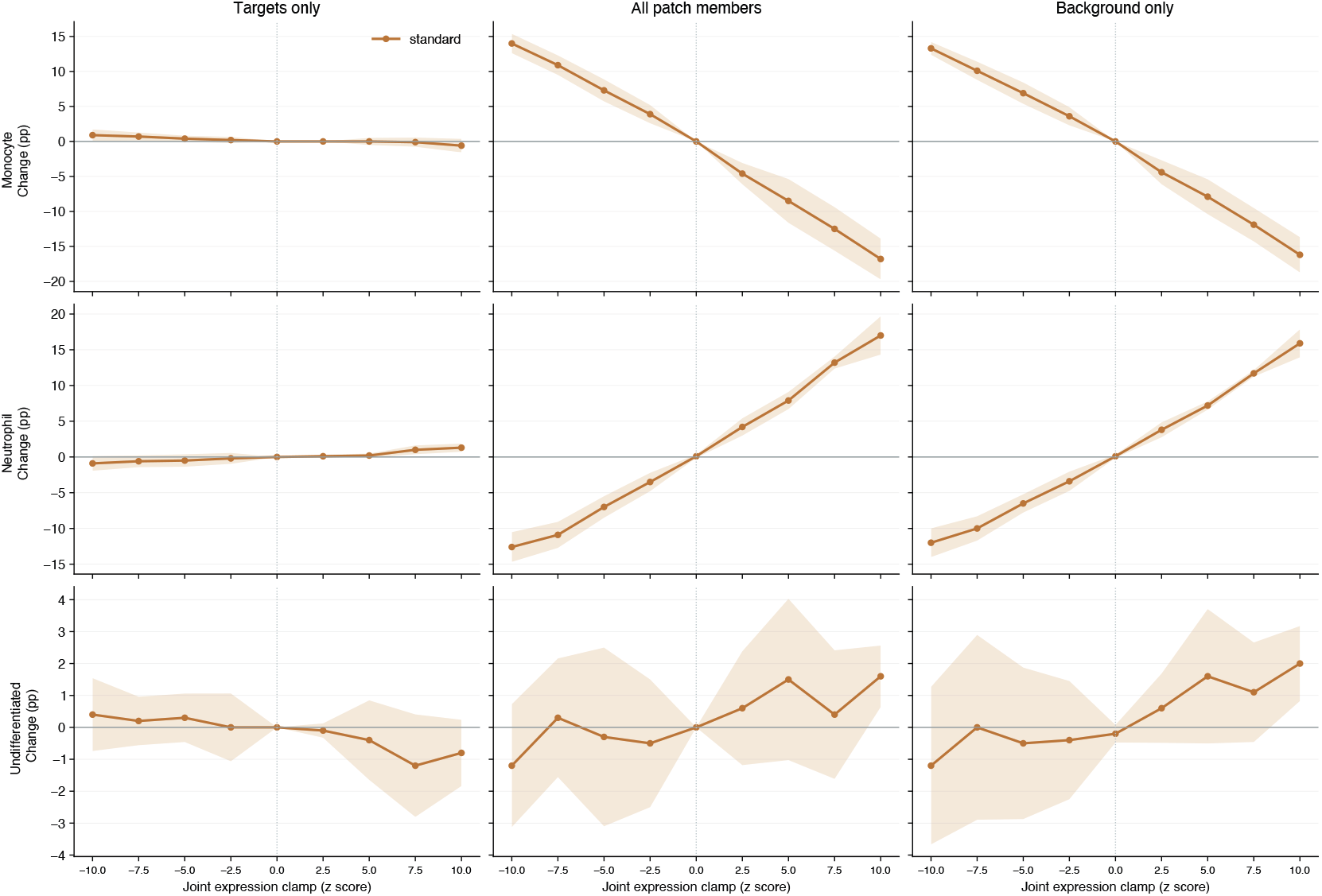
Geometric-context LARRY responses. Standard-prior seed 3; changes are relative to the untreated control. Columns are target only, all patch members, and background only—a different order from Figure S09. Background and whole-patch effects align and are larger than target-only effects. Bands are SD across five simulation groups. Different seeds and coupling settings limit causal comparison with Q2.

**Figure S11:**
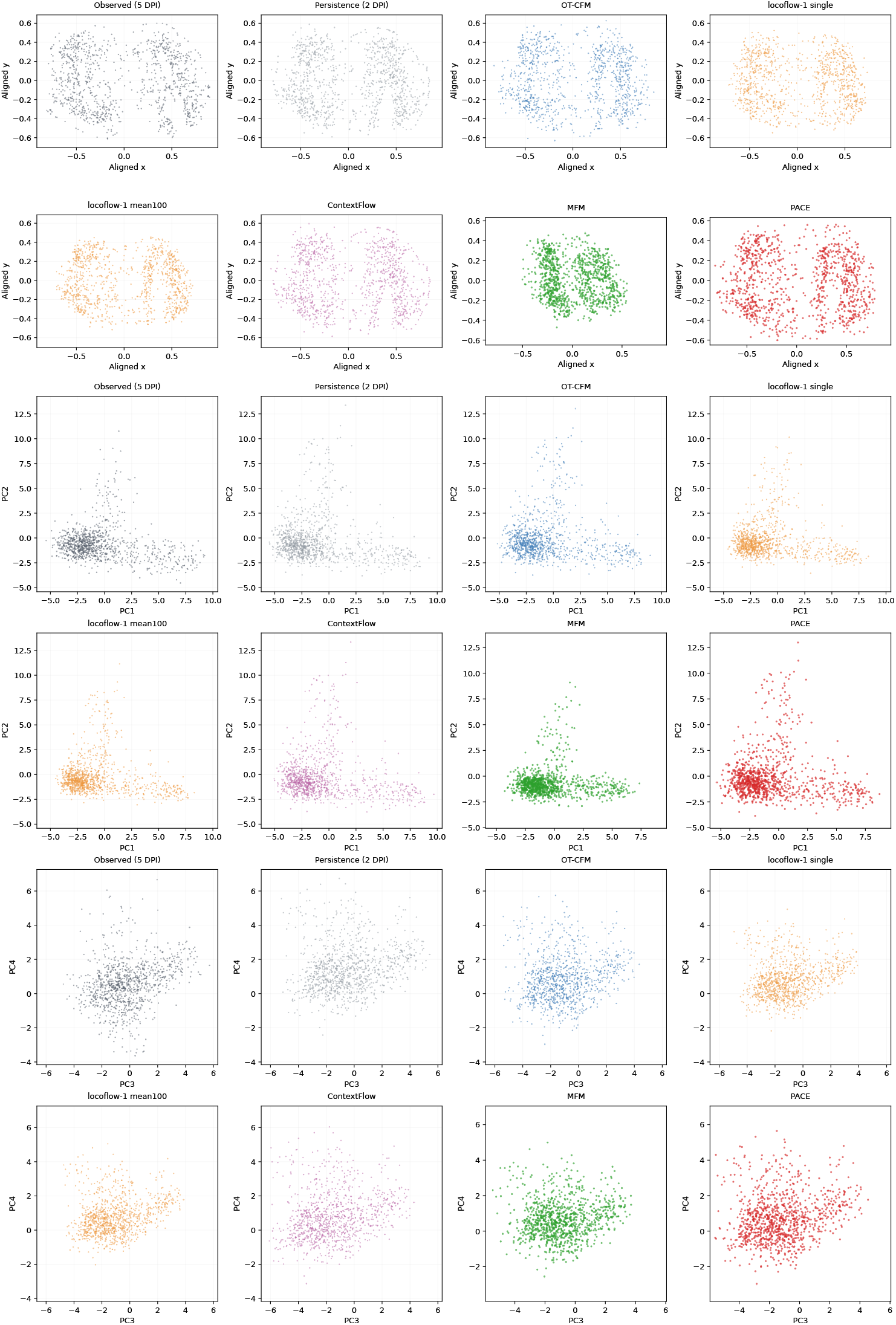
Spatial prediction examples. The top, middle, and bottom pairs of rows show measured coordinates, PC1/PC2, and PC3/PC4, respectively. Panels compare observed 5-DPI cells, persistence of 2-DPI cells, and model predictions; LOCOFLOW includes single and mean-100 predictions. Titles retain the supplied run and seed labels (0 or 1). These are individual-seed examples; Table 4 summarizes five training seeds. Axes are specific to each coordinate view.

**Figure S12:**
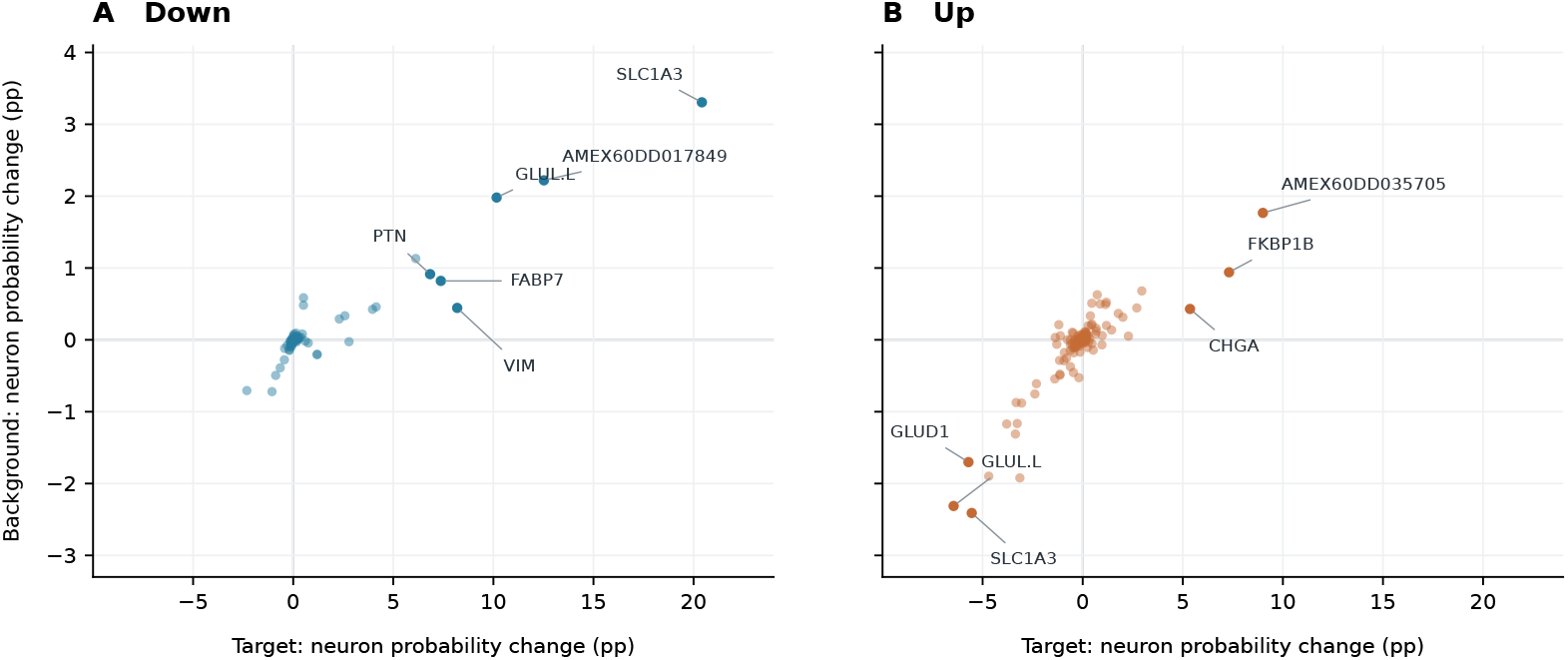
Predicted effects of target and background gene pertubations on neuronal fate probability.

## Notes

### Competing Interest Statement

The authors have declared no competing interest.

